# PHOTORELEASE OF OXYTOCIN *IN VIVO* USING PHOTOSWITCHABLE NANOVESICLES MODULATES HIPPOCAMPAL CIRCUIT DYNAMICS AND SOCIAL BEHAVIOR

**DOI:** 10.64898/2026.08.07.743279

**Authors:** Jaume Taura, Hassan Tajarenejad, Lailun Nahar, Hejian Xiong, Srijitjh Reddy Mudiganti, Yuanyuan Jiang, Krishna Sharmah Gautam, Samuel Achilefu, Xin Yu, Zhenpeng Qin, Paul A. Slesinger

## Abstract

Achieving precise spatiotemporal control over neuropeptide delivery *in vivo* remains a major challenge, as conventional approaches lack temporal resolution and control over release kinetics or have limitations for *in vivo* applications. Here, we demonstrate that photoswitchable azobenzene-containing lipid nanovesicles (“azosomes”) enable light-controlled release of neuropeptides *in vivo* in the brain of awake moving mice. Azosomes were infused into the hippocampus via optofluidic cannulas and activated using light stimulation in freely behaving mice. *In vivo* release kinetics were systematically characterized using calcein-loaded azosomes by varying light power, pulse duration, and post-infusion time. Oxytocin (OT)-loaded azosomes were used to assess bioactivity and receptor specificity using dual-color fiber photometry with the genetically encoded OT sensor (MTRIA_OT_), alongside pharmacological blockade with the oxytocin receptor antagonist OVTA (Ornithine-VasoTocin Analog). Azosomes enabled robust, repeatable, and light-dependent cargo release *in vivo* with tunable kinetics governed by stimulation parameters, with release efficiency controlled by light power and pulse duration. The system maintained functional stability for several hours post-infusion, with near-complete release achievable within a ∼2-hour window and minimal baseline leakage prior to stimulation. Controlled OT delivery produced rapid, receptor-specific increases in MTRIA_OT_ signals and modulated CA2 hippocampal circuit activity, reducing excitatory neuronal transient frequency and amplitude while altering social interaction dynamics, including decreased latency to initiate contact. These findings establish, for the first time, photoswitchable nanovesicles as a versatile platform for spatiotemporally precise delivery of neuropeptides *in vivo*, overcoming key limitations of existing delivery strategies and providing a broadly applicable framework for manipulating neuromodulatory signaling with high temporal precision.

## Introduction

Neuropeptides are important modulators of neuronal activity and behavior^1^. While there are over one hundred neuropeptides expressed in the brain^2,3^, our understanding of their functional role in brain circuitry is poorly understood. In general, neuropeptide signaling is considered slower (i.e., tens of seconds) and more diffuse than classical neurotransmission^4,5^. Recent studies using genetically encoded neuropeptide sensors, however, suggest that neuropeptide signaling can occur on relatively fast timescales (seconds), depending on the brain region and behavioral context^6–8^. While such sensors have enabled direct monitoring of neuropeptide dynamics *in vivo*, approaches to selectively and temporally control neuropeptide release remain limited. Optogenetic or chemogenetic activation of neuropeptide-producing neurons is one approach^9,10^; however, these neurons commonly co-release other neurotransmitters or neuropeptides making it difficult to establish a causal relationship between a specific neuropeptide, circuit activity, and behavior^1,11^.

Opto-pharmacological approaches have been developed for light-controlled release of transmitters. Caged compounds enable photo-activation with high temporal precision and have been successfully applied to both small molecules^12^ and neuropeptides^12,13^. However, caging requires chemical modification of each ligand with a photolabile protecting group and is therefore typically tailored to individual compounds, limiting its generalizability. Neuropeptides are subject to rapid peptidase degradation *in vivo*. In addition, caged molecules that are not fully inert prior to uncaging, particularly *in vivo*, may result in off-target biological activity^14^. Nanocarrier-based systems, including lipid vesicles, provide an alternative method for delivering bioactive molecules, but are primarily designed to enhance stability and retention, and often exhibit limited control over release kinetics once administered^15^. Together, these limitations highlight the need for a broadly applicable approach that enables externally controlled, temporally defined delivery of signaling molecules in the brain.

To address this gap in the field, we developed a technique of combining a photoswitchable lipid, azobenzene-phosphatidylcholine (AzoPC), with lipid nanovesicles (termed “azosomes”^16^) to provide a tool for light-controlled release of neuropeptides *in vivo*. Light-induced isomerization of Azo-PC in azosomes alters the permeability of lipid nanovesicle, enabling rapid release of their cargo on the timescale of milliseconds^16^. We showed previously that a dopamine D1 receptor agonist can be incorporated into azosomes and can activate cultures of striatal neurons *in vitro* upon photostimulation Here, we developed azosomes for controlled release of a neuropeptide *in vivo*, for the first time, and directly link neuropeptide release to circuit activity and behavior in awake, moving mice. We focus on the neuropeptide oxytocin (OT) for its well-known role in social behavior^17^ and show how OT influences endogenous signaling and social interactions. These results establish azosomes as a versatile platform for studying neuromodulatory systems *in vivo* with high temporal precision.

## Results

### Light-triggered release of cargo *in vivo* with azosomes

To establish the release of neuropeptides *in vivo* in awake, behaving mice, we first optimized the use of azosomes for light-dependent release of fluorescent dye in vivo. Azosomes were prepared by incorporating azobenzene-modified phosphatidylcholine (Azo-PC, 12 mol%) into lipid vesicles composed of DSPC, cholesterol, and Azo-PC at a molar ratio of 58:30:12 (see Methods), and encapsulating calcein dye. Irradiation at 365 nm switches the Azo-PC from cis to trans, increasing membrane permeability and allowing the cargo to permeate the liposome membrane^16^ (**Fig. 1A–B**). Cryo-TEM shows that azosomes are spherical membrane bilayers with an average diameter of ∼160 nm (**Fig. 1C**), a size that is expected to restrict diffusion of the azosomes in the brain. We first compared the effects of 365 nm photostimulation on calcein-loaded liposomes (i.e., no Azo-PC) infused into the left nucleus accumbens core (NAc) with those on calcein-loaded azosomes infused into the contralateral hemisphere (**Fig. 1D** and **E**). Using a dual fiber system to measure calcein fluorescence (**Fig. 1F**), we detected an increase in fluorescence in both hemispheres as the fibers approached the injection sites (**Fig. 1Gi**). After fiber stabilization (∼ 20 minutes), a brief 365 nm pulse (200 ms, 175 mW/cm²) was delivered that increased fluorescence in the azosome-containing hemisphere (green trace) but not in the calcein-filled liposome control (blue trace) (**Fig. 1Gii**). The increase in fluorescence arises from dequenching of calcein upon release^18^. Baseline fluorescence remained stable before stimulation, consistent with stable encapsulation of calcein (i.e., minimal leakage) *in vivo*. Two hours after implantation, a second 200 ms pulse of the same power produced only a modest increase in fluorescence (**Fig. 1Giii**), suggesting partial depletion of readily releasable cargo following the initial stimulation. However, increasing the stimulation duration to 2 s elicited a large increase in fluorescence (**Fig. 1Giv**). These results suggest that azosomes are stable for at least 2 h without leakage, and can support pulsatile stimulation and release. Finally, post hoc histological analysis confirmed placement in the NAc core (**Fig. 1H**) and little diffusion within the surrounding extracellular space. Together, these results demonstrate that azosomes enable stable, spatially confined, and repeatable light-triggered cargo release *in vivo*, with minimal leakage.

**Figure 1.**
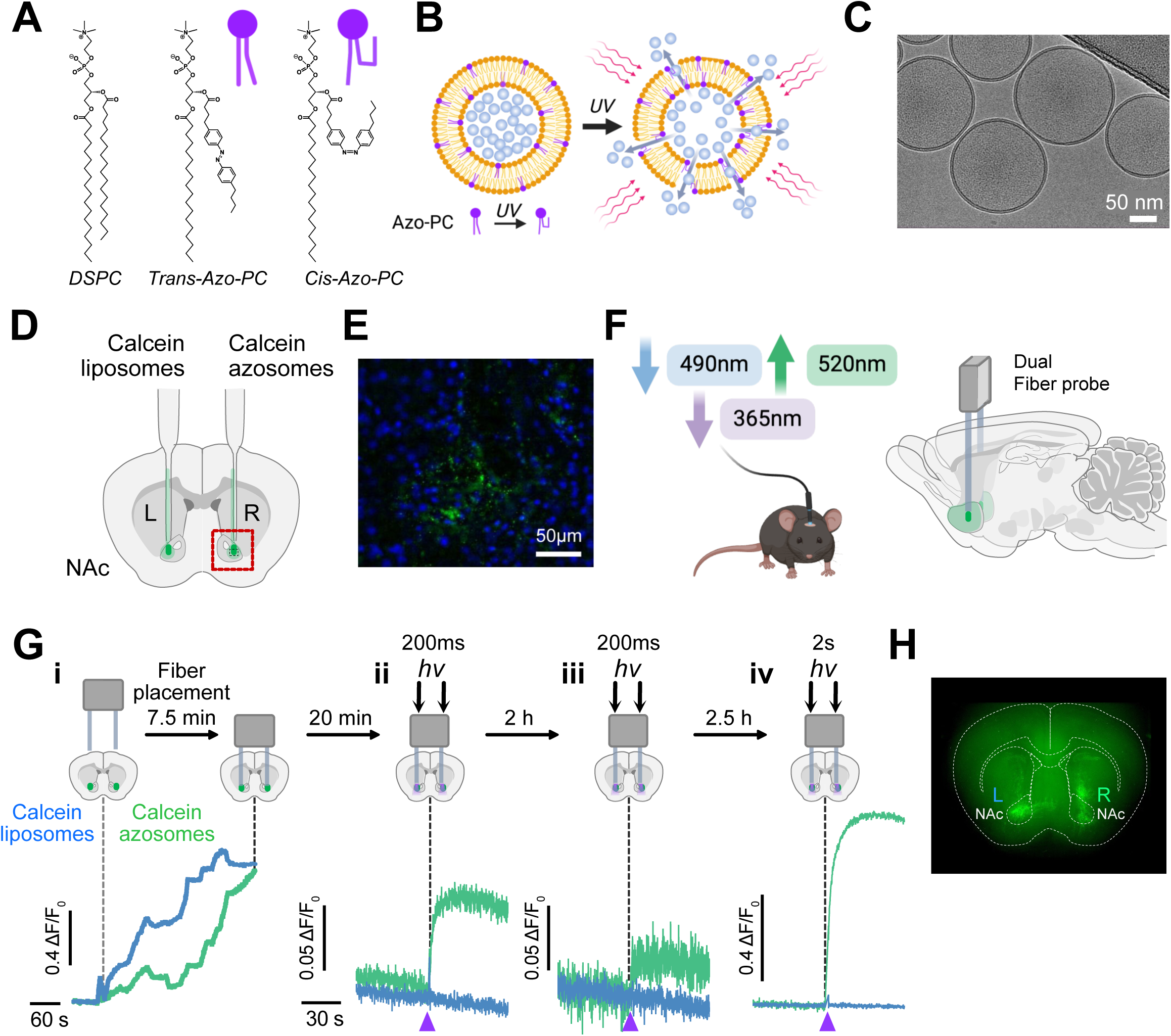
Light-triggered cargo release from photoswitchable nanovesicles *in vivo*. (**A**) Chemical structures of DSPC and azobenzene-modified phospholipid (Azo-PC), illustrating trans-to-cis isomerization upon 365 nm irradiation. (**B**) Schematic of azosome structure and light-induced increase in membrane permeability enabling cargo release. (**C**) Representative cryo-TEM image of azosomes showing spherical morphology (∼160 nm diameter). (**D**) Experimental design: bilateral infusion of calcein-loaded liposomes (left hemisphere) and calcein-loaded azosomes (right hemisphere) into the nucleus accumbens core (NAc). (**E**) Fluorescence image showing distribution of vesicles (green) within brain slice; nuclei labeled with DAPI (blue). (**F**) Schematic of fiber photometry system enabling simultaneous calcein excitation (490nm), calcein emission (520nm) and azobenzene isomerization (365 nm). (**G**) Dual fiber bilateral recordings and light delivery. *In vivo* fluorescence recordings during fiber approach (**i**) and light stimulation (**ii-iv**). (**ii**) Initial stimulation (200 ms, 175 mW/cm²) increases fluorescence in the azosome hemisphere but not in the liposome hemisphere. (**iii**) A second 200 ms pulse at 2 h produces a reduced response. (**iv**) A longer 2 s pulse reveals robust increase in fluorescence. Control liposomes show no response to 365 nm (**ii-iv**). (**H**) Post hoc histological verification of injection sites in the NAc.

To enable repeated delivery and systematic characterization of azosome-mediated photorelease *in vivo*, we implanted a commercially available optical fiber integrated with a cannula ), which provides an optical path and port for infusion (**Fig. 2Ai** and **2B**). Using the opto-fluidic cannula (OmFC) allows multiple infusions in the same animal over extended periods, enabling direct comparison of stimulation parameters within subjects. We first validated the opto-fluidic system with azosomes *in vitro* using a 0.5% agarose gel, which mimics the mechanical properties of brain tissue. Infusion of calcein-loaded azosomes into the agarose matrix resulted in a localized fluorescence signal corresponding to the distribution of vesicles (**Fig. 2Aii–iii, Supplementary Video 1**). Upon 365 nm irradiation, fluorescence increased initially near the fiber tip and rapidly spread throughout the infused region (**Fig. 2Aiv–v, Supplementary Video 1**), consistent with local release, dequenching, and diffusion of calcein. Quantification of fluorescence (ΔF/F_0_) confirmed a marked increase following a photostimulation pulse (**Fig. 2Avi**).

**Figure 2.**
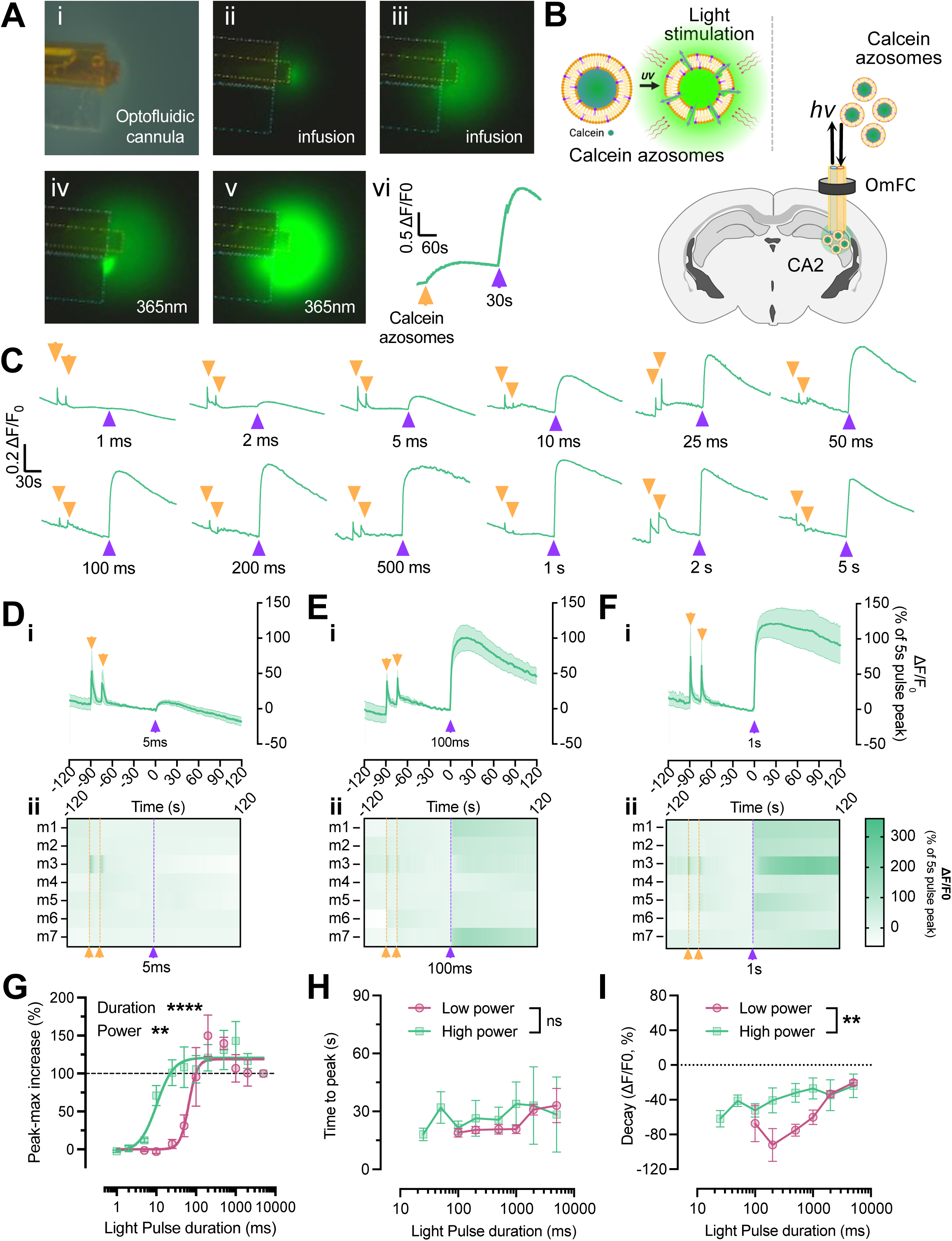
Opto-fluidic delivery enables repeated *in vivo* infusion and quantitative control of azosome photorelease. (**A**) *In vitro* validation of opto-fluidic cannula (OmFC) in 0.5% agarose. (**i**) Brightfield image of cannula positioned in agarose. (**ii–iii**) Fluorescence images during infusion of calcein-loaded azosomes. (**iv–v**) Fluorescence increase following 365 nm photostimulation, showing local release and spatial spread. (**vi**) Plot shows increase in fluorescence (ΔF/F) as a function of time (Supplementary Video 1). Orange triangles reflect time of injections. Purple triangles reflect time of 365 nm irradiation. (**B**) Schematic of *in vivo* opto-fluidic infusion and photometry recording in hippocampal CA2. (**C**) Representative *in vivo* fluorescence traces showing responses across light pulse durations (1 ms to 5 s; high power condition). (**D-F**) Averaged *in vivo* fluorescence responses (**i**, mean ± SEM) and corresponding heatmaps (**ii**) for pulse durations of 5 ms, 100 ms, and 1 s. (**G**) Peak fluorescence increase as a function of light pulse duration at low and high power. Curves fitted using standard 4 parameter Hill plot. (**H**) Time to peak fluorescence across stimulation conditions is not different. (**I**) Signal decay, calculated as the percentage decrease in ΔF/F between 25 s and 115 s post-stimulation, showing less decay at higher power and longer pulse durations. Data were analyzed using two-way ANOVA with factors power and pulse duration. *p < 0.05, **p < 0.01, ***p < 0.001, ****p < 0.0001. Data are presented as mean ± SEM (n = 4–10).

We next examined the *in vivo* properties of dye-filled azosomes in the hippocampal CA2 region (**Fig. 2B**). Following recovery (2-3 weeks) from implantation of the opto-fluidic cannula , calcein-loaded azosomes were infused and then photostimulated while measuring fluorescence using fiber photometry (**Fig. 1F and 2B**). Real-time recordings showed that each infusion produced a small transient increase in fluorescence with the dye-filled azosomes (**Fig. 2C, yellow triangle**). We examined the effect of increasing duration of photostimulation pulses (1 ms to 5 s) at both low (175 mW/cm²) and high power (1750 mW/cm²). Short pulses (2–5 ms) produced detectable release of calcein (increase in ΔF/F_0_), while intermediate durations (25–100 ms) elicited larger, transient responses, and longer pulses (0.5–1 s) resulted in maximal and sustained fluorescence increases. Quantification of the fluorescent response following 5 ms, 100 ms and 1 s photostimulation pulses (**Fig. 2D-F**, top, **i**) shows that longer pulses producing both higher peak fluorescence and sustained signals. The heat map plots show a consistent change in all seven mice (**Fig. 2D-F**, bottom, **ii**). We plotted the peak fluorescence (**Fig. 2G**) as a function of pulse duration for the two different stimulation powers, and found significant effects of both duration (two-way ANOVA, p < 0.0001) and power (p = 0.0066), with higher power shifting the response toward shorter durations. In contrast, the time to peak fluorescence was not significantly affected by either duration or power (**Fig. 2H**), remaining approximately constant (∼30 s), suggesting that calcein dequenching is rapid, and that diffusion of calcein underlies the decay. Consistent with this, analysis of the signal decay (**Fig. 2I**) showed that higher power and longer pulse durations produced more sustained fluorescence signals (duration: p = 0.0152; power: p = 0.0016), consistent with greater dye release. Further support for this interpretation comes from the area under the curve (AUC) analysis (**Supplementary Fig. S1**), which shows a saturation of the response at longer pulse durations, indicating that extended stimulation increases total release rather than peak amplitude. Together, these results demonstrate that azosome-mediated photorelease can be finely tuned *in vivo* by adjusting light stimulation parameters, with rapid release achieved within tens of milliseconds, and maximal release efficiency and signal persistence obtained with higher power and stimulation durations on the order of seconds.

### Azosomes remain functionally stable *in vivo* for at least 1 hour

To assess the stability of azosomes *in vivo* and their ability to retain photo-responsive properties over time, we conducted an *in vivo* time course. We used photostimulation parameters that maximized release (1 s, 365 nm, high power) and examined the photorelease of azosomes after varying amounts of time (Δt = 1 min to 14 h) following infusion *in vivo* (**Fig. 3A**). Representative fluorescence traces (**Fig. 3B**) show that robust photo-release of calcein was maintained for up to 1 h following infusion, with comparable peak responses across early time points. With longer delays, photorelease efficiency progressively declined, although detectable responses were still observed several hours after infusion. Quantification of peak fluorescence responses (**Fig. 3C**) revealed that azosome photorelease remained largely stable within the first 1–2 h, followed by a gradual reduction in efficiency, with approximately 50% decrease observed at 3–4 h and near-complete loss by 14 h. Fitting the decay with a one-phase exponential model yielded a half-life of ∼187 min, indicating a slow loss of functional capacity *in vivo*. Taken together, these findings indicate that azosomes retain their cargo and photo-responsiveness in the brain parenchyma for extended periods following infusion, suitable for photo-releasing during a behavioral paradigm.

**Figure 3.**
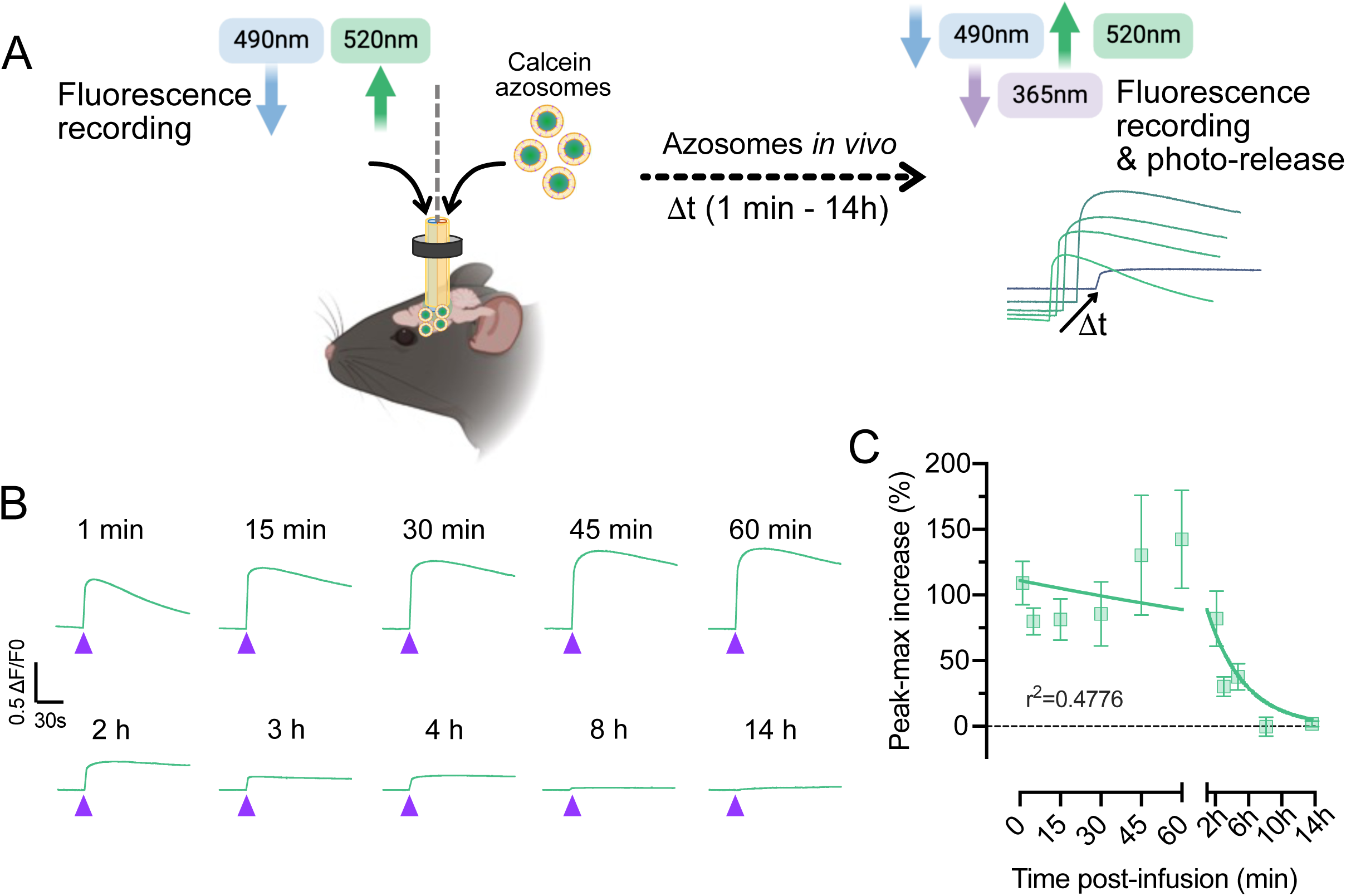
Azosomes remain stable *in vivo* over extended time periods. (**A**) Schematic shows assessment of azosome stability *in vivo*. Calcein-loaded azosomes were infused through the opto-fluidic cannula, and photorelease was triggered at defined time points ranging from 1 min to 14 h post-infusion using 1 s pulses of 365 nm light at high power. Fluorescence was recorded via fiber photometry. (**B**) Representative fluorescence traces showing light-triggered release at different time points following infusion. Robust photorelease is observed up to 1 h, with progressive reduction in response at longer intervals. (**C**) Quantification of peak fluorescence increase (ΔF/F, normalized to maximal response) as a function of time post-infusion. Data show stable photorelease within the first 1–2 h, followed by a gradual decline in efficiency over time. Data were fitted using a one-phase exponential decay model. Peak responses were fitted using a one-phase decay model (R² = 0.4776; half-life = 186.9 min).

### Validation of *in vivo* oxytocin detection using MTRIAOT

We next tested whether azosomes could be used to encapsulate and photorelease a neuropeptide *in vivo*. We focused on oxytocin (OT) because of its well known role in modulating social behaviors^17^ and because it can be detected with a genetically encoded sensor MTRIA_OT_^19^. An opto-fluidic cannula was implanted into the dorsomedial striatum (DMS) of mice, and after 2-3 weeks, AAV encoding MTRIA_OT_ was infused through the cannula. After 3–4 weeks, fluorescence signals were recorded using a three-color fiber photometry system allowing simultaneous monitoring of MTRIA_OT_ activity and infusions (Calcein Red 10µM) (**Fig. 4A**). To characterize MTRIA_OT_ responses *in vivo*, we examined the effect of infusing increasing concentrations of OT (10 nM to 100 μM) alongside vehicle controls; Calcein Red was mixed with the azosomes to monitor solution delivery. Representative traces (**Fig. 4B**) show successful infusion across all conditions (red trace), whereas consistent MTRIA_OT_ responses (green trace) were observed at OT concentrations of 1 µM or higher. Quantification revealed a clear dose-dependent response, with significant increases at 10 μM and 100 μM (**Fig. 4C,D**; one-way ANOVA, p < 0.0001). To validate the specificity of the MTRIA_OT_ signal, we examined the effect of the oxytocin receptor (OXTR) antagonist OVTA. Sequential infusions within the same session showed that OT (10 μM) induced a robust increase in MTRIA_OT_ fluorescence, whereas OVTA (100 μM) produced an opposite (negative) response, consistent with competitive displacement of OT at the MTRIA_OT_ receptor. Importantly, co-application of OT with OVTA abolished the OT-induced MTRIA_OT_ signal (**Fig. 4E,F**). Together, these results demonstrate that the fluorescence changes for MTRIA_OT_ are dependent on OT, and that azosome-based platforms can be paired with genetically encoded sensors to monitor neuropeptide signaling with high specificity *in vivo*.

**Figure 4.**
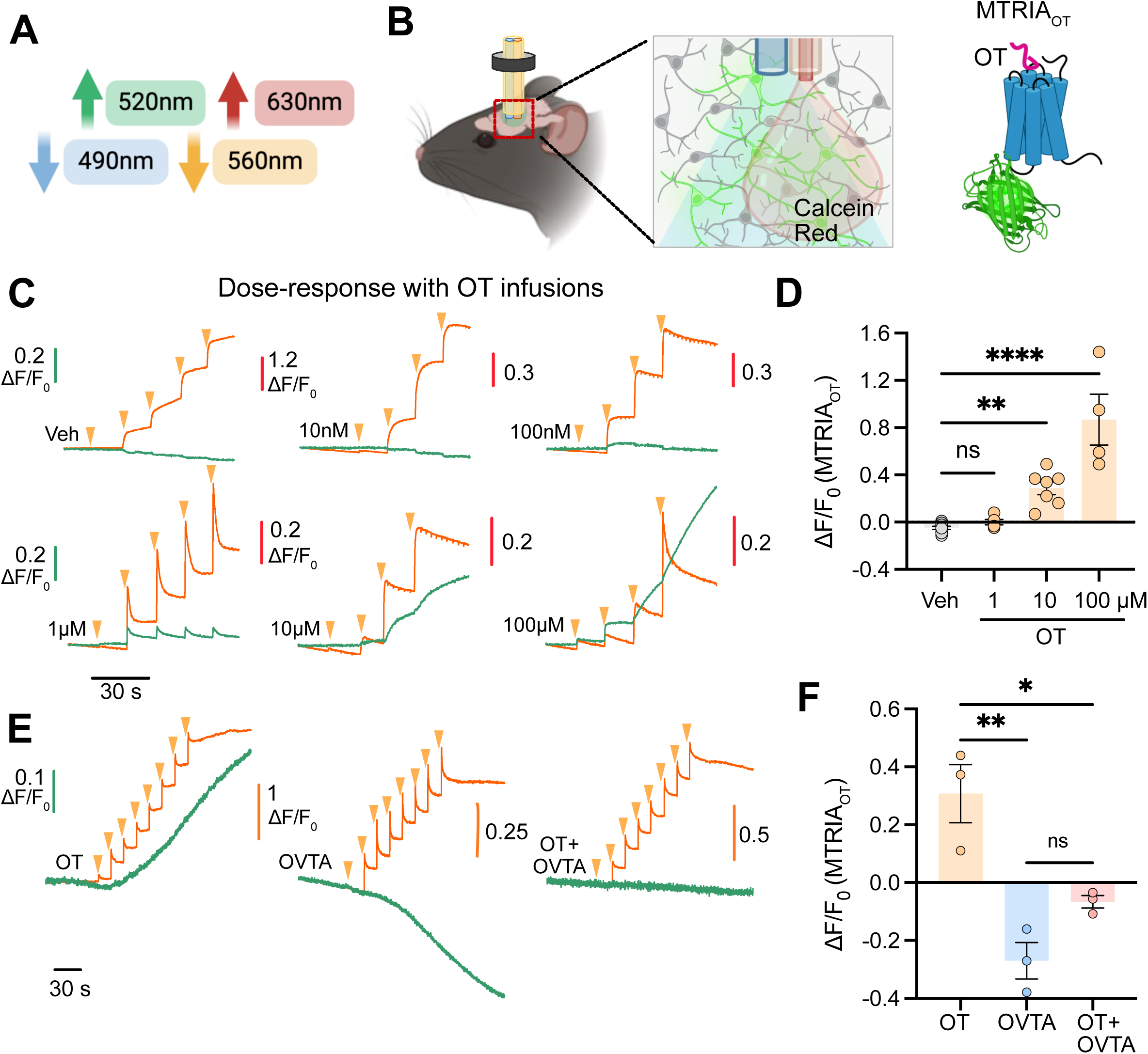
Detection of oxytocin *in vivo* using opto-fluidic delivery and MTRIA_OT_ sensor. (**A**) Schematic of the fiber photometry system enabling simultaneous detection of MTRIA_OT_ (490nm Ex; 520nm Em) and Calcein Red (560nm Ex; 630nm Em). (**B**) The opto-fluidic cannula allows combined optical recording and local infusion of MTRIA_OT_ virus as well as other compounds into the dorsomedial striatum (DMS). (**C**) Representative fluorescence traces showing infusion of vehicle compared to increasing concentrations of free oxytocin (10 nM, 100 nM, 1 μM, 10 μM, 100 μM). Calcein Red fluorescence allows monitoring of solution delivery – note ‘staircase’ increase for each injection. MTRIA_OT_ fluorescence reflects detection of oxytocin. (**D**) Quantification of MTRIA_OT_ responses (ΔF/F₀) across oxytocin concentrations (n = 4–10). Significant responses were observed at 10 μM and 100 μM, indicating dose-dependent sensor activation. One-way ANOVA: F(3,22) = 27.60, p < 0.0001. Tukey’s multiple comparisons: Vehicle vs OT-10 μM (**p = 0.0049), Vehicle vs OT-100 μM (****p < 0.0001), OT-10 μM vs OT-100 μM (***p = 0.0002); OT-1 μM ns. (**E**) Representative traces show effect of OT receptor antagonist (OVTA) on MTRIA_OT_. Sequential infusions of first OT (10 μM), then OVTA (100 μM) alone, and then OT (10 μM) + OVTA (100 μM) OT + OVTA. (**F**) Quantification of MTRIA_OT_ responses for OT (10 μM), OVTA (100 μM) alone, and OT (10 μM) + OVTA (100 μM) (n = 3). OVTA alone reduces basal fluorescence and antagonizes OT-induced responses. (**E**) One-way ANOVA: F(2,6) = 17.72, p = 0.0030. Tukey’s test: OT vs OVTA (**p = 0.0026), OT vs OT+OVTA (*p = 0.0209), OVTA vs OT+OVTA ns.

### Azosome-mediated photorelease of oxytocin *in vivo*

To determine whether azosomes enable controlled release of OT *in vivo*, we next tested OT-loaded azosomes with MTRIA_OT_. We first measured the stability of azosomes and found that OT azosomes are stable at 37 C for at least 2h (**Supplementary Fig. S2A,B**). OT-azosomes were infused into the DMS of a mouse expressing MTRIA_OT_ while simultaneously monitoring injection via Calcein Red fluorescence and OT signaling via MTRIA_OT_ (**Fig. 5A**). Following infusion, the field was irradiated with a 1 s light pulse (365 nm, high power) to trigger neuropeptide release. A representative recording shows that vehicle infusion produced a clear increase in red fluorescence without affecting the MTRIA_OT_ signal (**Fig. 5B, top**). Subsequent irradiation produced some acute photobleaching but no persistent increase in fluorescence, indicating little effect of irradiation on MTRIA_OT_. In contrast, infusion of OT-azosomes produced minimal changes in MTRIA_OT_ fluorescence, likely due to a low amount of free OT (**Supplementary Fig. S2C**) but a marked increase in MTRIA_OT_ signal following irradiation (**Fig. 5B, middle**). As a control, OT (10 μM) delivered at the end of the recording elicited a robust increase in MTRIA_OT_ fluorescence during infusion, with no additional effect following irradiation (**Fig. 5B, bottom**). Quantification of the ΔF/F_0_ across animals confirmed these observations (**Fig. 5C,D**). During infusion, only OT (10 μM) induced a significant increase in MTRIA_OT_ signal, whereas OT-azosomes without irradiation did not differ from vehicle (one-way ANOVA, p < 0.0001) (**Fig. 5C**). Remarkably, OT-azosomes produced a significant increase in MTRIA_OT_ signal following irradiation, while vehicle and OT conditions did not show light-dependent effects (p = 0.0010) (**Fig. 5D**). The magnitude of the response elicited by OT-azosome photorelease was comparable to that observed with direct OT infusion. Similarly, photo-release of OT from azosomes *in vitro* released approximately 25 µM OT following a 1 s pulse (**Supplementary Fig. S2C**). To further assess receptor specificity, we performed sequential infusion–irradiation experiments in the presence of the OXTR antagonist OVTA. An initial irradiation of OT-azosomes produced the expected increase in MTRIA_OT_ fluorescence (**Fig. 5E, top**). Infusion of OVTA then resulted in a decrease in MTRIA_OT_ fluorescence (as shown above), consistent with displacement of bound OT (**Fig. 5E, middle**). Under these conditions, subsequent irradiation of OT-azosomes failed to induce an increase in MTRIA_OT_ signal, in contrast to control (**Fig. 5E, top**). Quantification of the ΔF/F_0_ showed that OVTA significantly reduced MTRIA_OT_ signal during infusion and abolished the irradiation-induced response observed with OT-azosomes (one-way ANOVA, p = 0.0193 for infusion; p < 0.0001 for irradiation) (**Fig. 5F,G**). Together, these results demonstrate that azosomes enable photo-release of oxytocin *in vivo*, producing receptor-dependent signaling that can be controlled temporally and monitored simultaneously using genetically encoded sensors.

**Figure 5.**
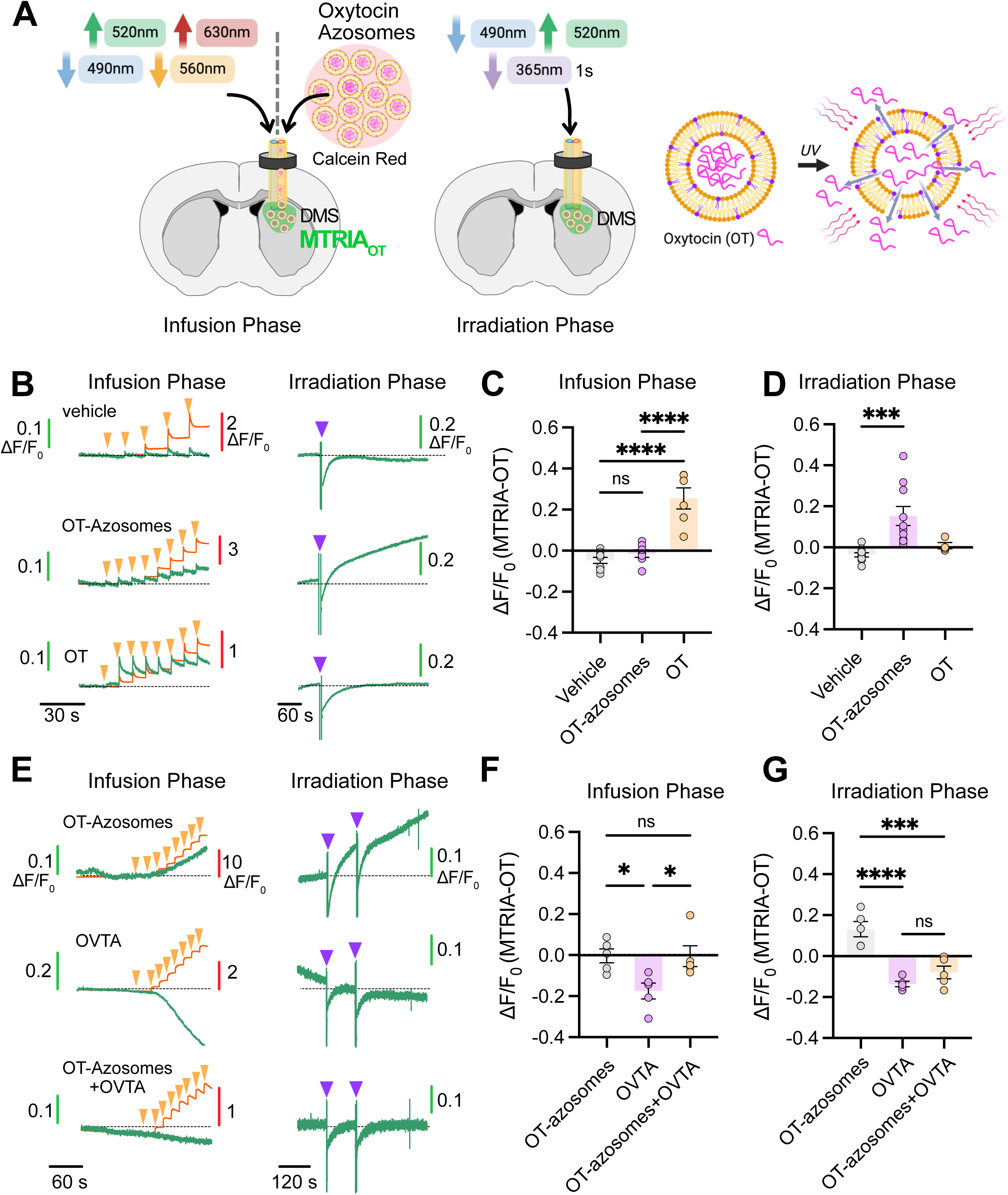
**Light-triggered release of OT from azosomes *in vivo.*** (**A**) Schematic of the experimental design combining opto-fluidic delivery and fluorescence recording in the DMS. OT-loaded azosomes are infused while monitoring MTRIA_OT_ (490nm Ex; 520nm Em) and Calcein Red (560nm Ex; 630nm Em) signals. Following infusion, local irradiation (365 nm) is applied to trigger oxytocin release from azosomes. (**B**) Representative fluorescence traces from a single animal showing sequential infusion and irradiation of vehicle, OT-azosomes, and OT (1 μM). Calcein Red fluorescence reports solution delivery (4-8 injections of x nl each; see Methods). Top traces: MTRIA_OT_ signal does not change during infusion or irradiation for vehicle. Middle traces; MTRIA_OT_ signals increase slightly during OT-azosomes infusion and robustly following irradiation. Bottom traces; MTRIA_OT_ signals increase during OT infusion but not following irradiation. Transient signal deflection during irradiation reflects optical artifact. (**C,D**) Quantification of MTRIA_OT_ responses (ΔF/F₀) during infusion (**C**) and irradiation (**D**) across conditions (n = 6–10). OT (10 μM) induces significant responses during infusion, whereas OT-azosomes elicit significant responses only upon irradiation. One-way ANOVA: F(2,21) = 9.824, p = 0.0010. Tukey’s test: Vehicle vs OT-azosomes (***p = 0.0008), Vehicle vs OT-10 μM ns, OT-azosomes vs OT-10 μM ns. One-way ANOVA: F(2,19) = 32.04, p < 0.0001. Tukey’s test: Vehicle vs OT-10 μM (****p < 0.0001), OT-azosomes vs OT-10 μM (****p < 0.0001), Vehicle vs OT-azosomes ns. (**E**) Representative traces showing sequential infusion–irradiation experiments with OT-azosomes (Top traces) and OVTA (100 μM) (Middle traces) and OT-Azosomes plus OVTA (100 μM) (Bottom traces). OVTA induces a reduction in MTRIA_OT_ signal and prevents subsequent responses to OT-azosome photorelease. (**F,G**) Quantification of MTRIA_OT_ responses (ΔF/F₀) during infusion (**F**) and irradiation (**G**) for OT-azosomes, OVTA, and OT-azosomes + OVTA (n = 5). OVTA reduces baseline signal and abolishes irradiation-induced responses. One-way ANOVA: F(2,12) = 5.583, p = 0.0193. Tukey’s test: OT-azosomes vs OVTA (*p = 0.0325), OVTA vs OT-azosomes+OVTA (*p = 0.0345), OT-azosomes vs OT-azosomes+OVTA ns. One-way ANOVA: F(2,12) = 23.85, p < 0.0001. Tukey’s test: OT-azosomes vs OVTA (****p < 0.0001), OT-azosomes vs OT-azosomes+OVTA (***p = 0.0006), OVTA vs OT-azosomes+OVTA ns.

### Light-triggered oxytocin release modulates social behavior and endogenous signaling dynamics

Having established that we can photorelease OT in a spatially and temporally controlled manner *in vivo*, we next examined the biological effect of OT release on social behavior. We targeted the hippocampal CA2 region, which has high OXTR expression and is involved in social behaviors^20^, and combined opto-fluidic delivery with *in vivo* while monitoring MTRIA_OT_ during a social interaction task. Male mice were implanted with opto-fluidic cannulas and subsequently infused with AAV encoding MTRIA_OT_ (**Fig. 6A**). OT-azosomes were infused under anesthesia, and then mice were allowed to recover. Mice were then transferred to a behavioral arena in which social interaction was initiated by opening a gate that allowed access to a compartment containing a female mouse enclosed in a cup. In alternating sessions, animals either remained in dark conditions or received a brief light pulse prior to social interaction to trigger OT release (**Fig. 6A**). Fluorescence recordings during the infusion phase showed comparable delivery across conditions, as indicated by similar Calcein Red signals, with no significant increase in MTRIA_OT_ during infusion (**Fig. 6B**). In contrast, irradiation induced a clear increase in MTRIA_OT_ fluorescence that remained elevated for several minutes, indicating increased OT during the subsequent social interaction period (**Fig. 6B**). To characterize behavioral responses, social interactions were quantified using a combination of position tracking and a capacitive sensor integrated into the social stimulus enclosure (**Fig. 6A,C and Supplementary Fig. S5**, see methods). This approach enabled precise, subsecond detection of social-contact events based on gate status, proximity, and physical interaction (**Fig. 6C,D**). Quantification of social behavior revealed that the number of social-contact events did not differ between conditions (**Fig. 6Ei**; paired t-test, p = 0.3363). However, the duration of social contacts was significantly reduced with photo-released OT, compared to control (**Fig. 6Eii**; p = 0.0334). The latency to first interaction was also significantly decreased (p = 0.0171), indicating faster initiation of social behavior following OT photorelease (**Fig. 6Eiii**). We next examined MTRIA_OT_ dynamics aligned to social-contact events. In control conditions, endogenous OT signaling exhibited a temporal pattern characterized by a transient decrease preceding contact, followed by a post-contact increase peaking within ∼5–15 s (**Fig. 6F,G**). In contrast, this temporal signature was altered following OT photorelease, with attenuation of both the pre-contact decrease and post-contact increase (**Fig. 6F,G**). Quantification showed significant reduction in the pre-contact trough amplitude (**Fig 6H,I**; unpaired t-test, p < 0.0001) and in the post-contact peak response (p = 0.0073) in the photo-release condition, as compared to control (no photostimulation). Together, these results demonstrate that azosome-mediated photorelease of oxytocin enables temporally controlled modulation of endogenous oxytocin signaling and alters social interaction dynamics *in vivo*.

**Figure 6.**
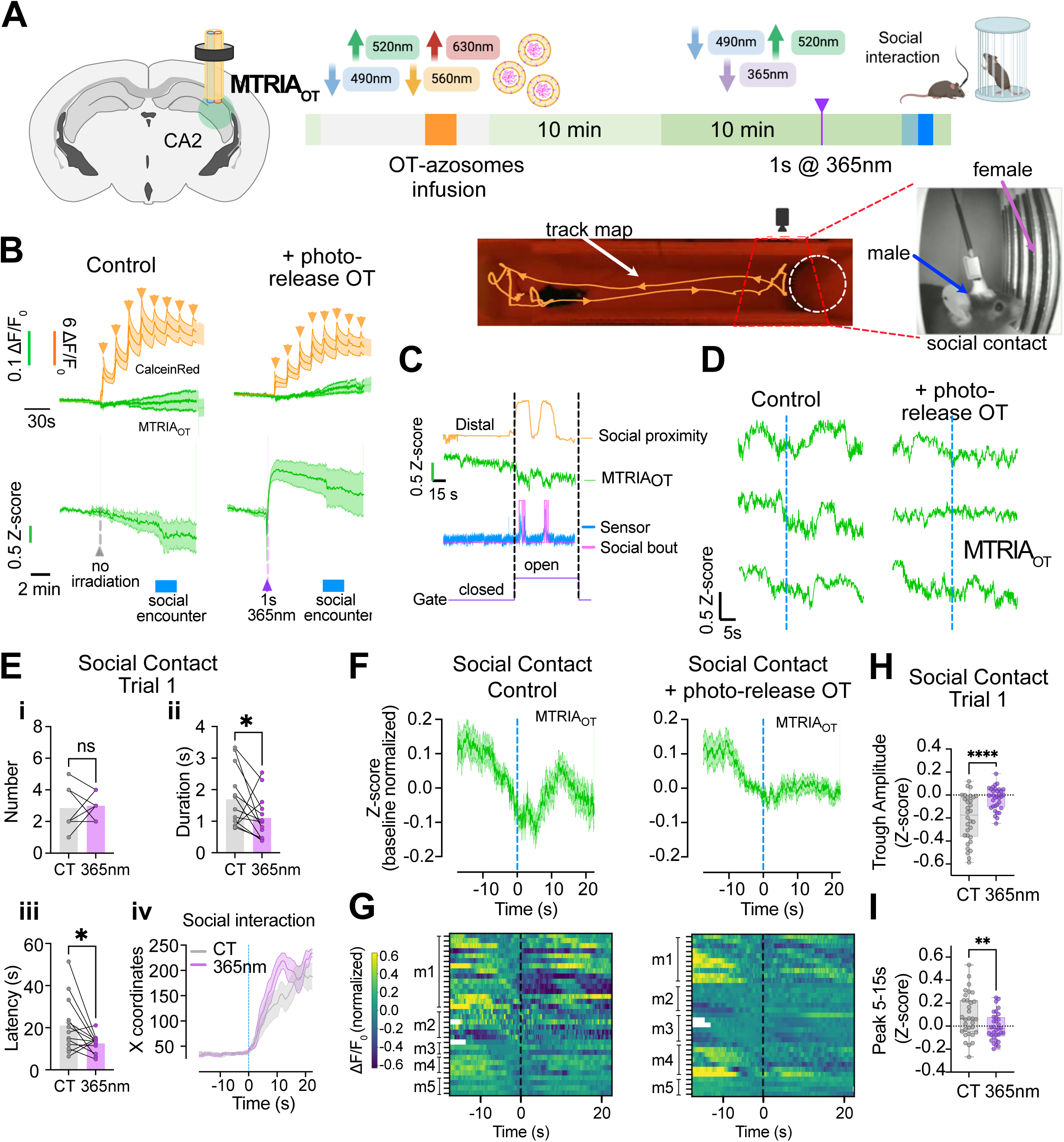
**Light-triggered OT release modulates social behavior and endogenous OT signaling dynamics**. (**A**) Schematic of opto-fluidic cannula implantation in the hippocampal CA2 region and expression of the MTRIA_OT_ sensor. Experimental timeline showing OT-azosome infusion, recovery, optional light stimulation (365 nm), and subsequent social interaction assay. Bottom, track map shows example of male mouse that pursues female in the cup. (**B**) Average (mean ± SEM, n= x) fluorescence signals for two groups during infusion (top traces) and post-infusion period (bottom traces) for control (no irradiation) and light (365 nm) conditions. Calcein Red allows tracking of OT-azosome injections. MTRIA_OT_ increases following irradiation and remains elevated during the behavioral session, in contrast to control (bottom traces). (**C**) Representative traces showing MTRIA_OT_ signal (green), gate status (purple), proximity (yellow), and capacitive sensor output (blue) used to define social-contact events. (**D**) Representative traces of MTRIA_OT_ aligned to individual social-contact events (dashed line) for control and light conditions. (**E**) Quantification of social behavior metrics, including number of social-contact events (**i**), contact duration (**ii**), and latency to first interaction (**iii**) (n = 13). Light-triggered OT release reduces contact duration and latency without affecting event number. Paired t-tests: number of contacts (p = 0.3363, ns), duration (*p = 0.0334), latency (*p = 0.0171). (**iv**) A plot of the position of the mouse versus time for the two conditions. (**F,G**) Averaged MTRIA_OT_ Z-score aligned to social-contact events for control and light conditions, with corresponding heatmaps for individual mice (m1-m5) (**G**). (**H,I**) Quantification of pre-contact trough amplitude (**H**) and post-contact peak (5–15 s) (**I**). Unpaired t-tests: trough amplitude (****p < 0.0001), peak 5–15 s (**p = 0.0073).

### Photorelease of oxytocin modulates CA2 glutamatergic activity during social interaction

To determine whether OT photorelease alters local circuit activity, we monitored calcium dynamics in CA2 glutamatergic neurons using Cre-dependent jGCaMP7s expression in Vglut1-Cre mice combined with opto-fluidic delivery of OT-azosomes (**Fig. 7A**). The behavioral paradigm and analysis framework were identical to those described previously (**Fig. 6A)**. Event-aligned analysis of vGLUT1-GCaMP signals revealed a structured temporal pattern around social-contact onset in control sessions, characterized by a transient decrease preceding contact followed by a delayed increase peaking ∼5–15 s after interaction (**Fig. 7B,C**). This pattern was markedly altered following OT photorelease, with attenuation of both pre-contact suppression and post-contact activation (**Fig. 7B,C**). A similar temporal profile was observed for MTRIA_OT_ signals. Given the fast and transient nature of calcium signals, we next quantified activity using peak-based analysis. Representative traces illustrate discrete calcium transients and their detection across conditions (**Fig. 7D**). Interestingly, raster plots aligned to social-contact events revealed a clear reduction in transient occurrence in the photo-release condition (**Fig. 7E**). Quantification confirmed a significant decrease in transient frequency following OT photorelease (**Fig. 7F**; Control: 0.483 ± 0.045 vs. 365 nm: 0.247 ± 0.025; unpaired t-test, p < 0.0001) as well as reduced transient amplitude (prominence) (Dark: 0.321 ± 0.008 vs. 365 nm: 0.249 ± 0.013; p < 0.0001) (**Fig. 7G**). Together, these results indicate that acute OT release suppresses CA2 glutamatergic activity during social interaction.

**Figure 7.**
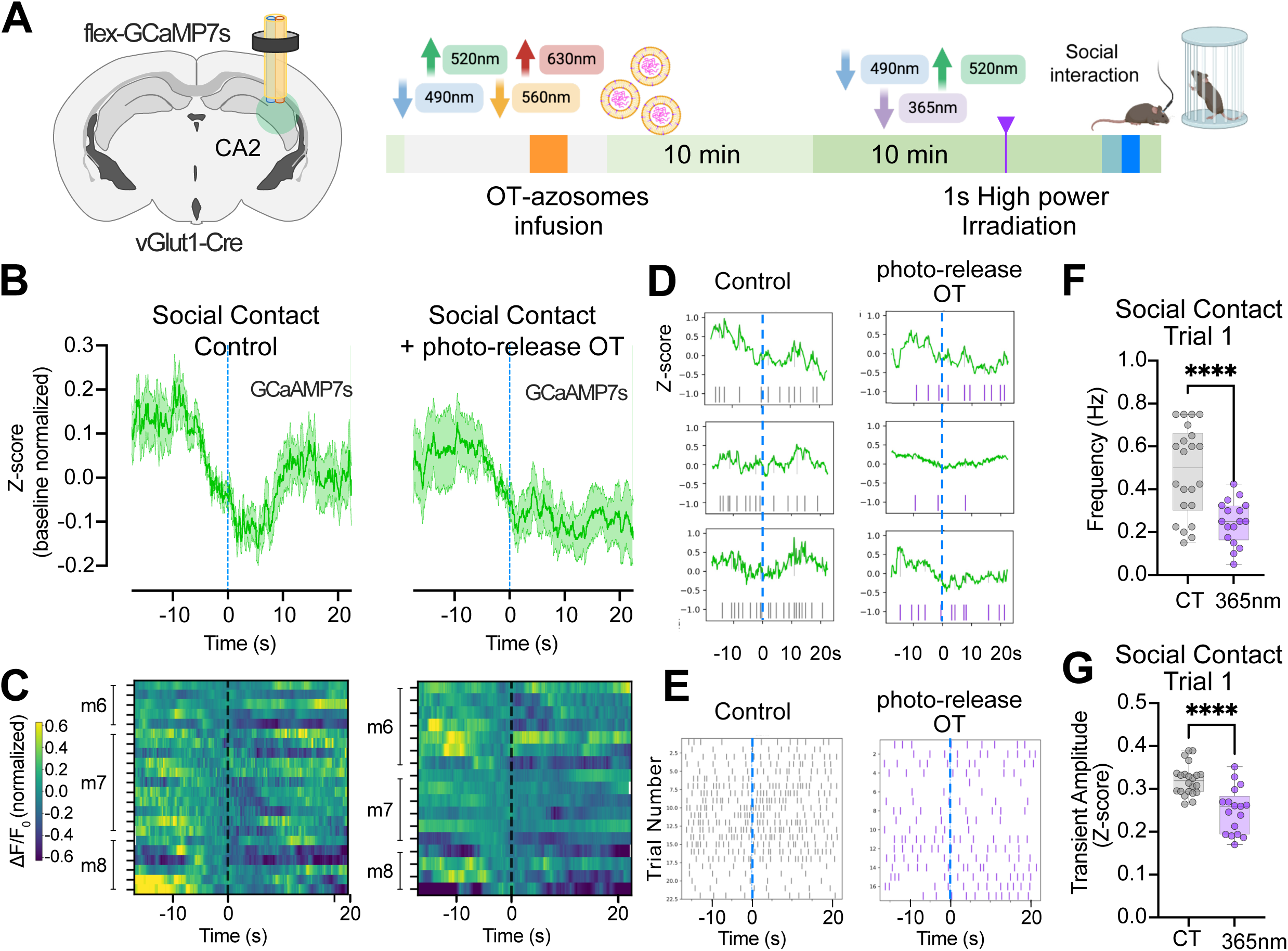
Photorelease of OT modulates CA2 glutamatergic activity during social interaction. (**A**) GCaMP7S expressed in glutamate neurons (VGlut1-Cre). Experimental timeline show OT-azosome infusion, recovery, optional light stimulation (365 nm), and subsequent social interaction assay. (**B,C**) Averaged MTRIA_OT_ Z-score aligned to social-contact events for control and light conditions, with corresponding heatmaps for individual mice (m6-m8) (**C**). Control sessions exhibit a biphasic response with pre-contact suppression and post-contact activation, which is attenuated following OT photorelease. (**D**) Representative vGLUT1-GCaMP traces during individual social-contact events in control and light conditions. Detected calcium transients are indicated by a line along each trace. (**E**) Raster plots of calcium transient occurrence aligned to social-contact onset across events and animals for each condition. Reduced transient density is observed following OT photorelease. (**F,G**) Quantification of calcium transient frequency (**F**) and amplitude (prominence) (**G**) during social-contact events. OT photorelease significantly reduces both transient frequency and amplitude (unpaired t-tests; ****p < 0.0001).

## Discussion

Understanding how neuropeptide signaling shapes brain circuits and behavior requires tools that enable precise spatiotemporal control over molecular delivery *in vivo*. Conventional approaches, such as local infusion through implanted cannulas^21,22^, allow targeted administration of compounds but lack temporal control. In this study, we introduce azosomes, containing a photoswitchable liposomes, as a platform for spatially and temporally controlled release of neuropeptides *in vivo* and demonstrate their ability to causally link release to circuit activity and behavior. By combining opto-fluidic delivery with fiber photometry, we show that light-triggered release of encapsulated neuropeptide can be precisely timed and monitored in freely behaving animals.

The implementation of azosomes for *in vivo* applications required consideration of several practical aspects. To establish their utility for neuropeptide photo-release *in vivo*, we first characterized and validated the performance of fluorescent dye-filled azosomes. These validation steps are likely to be important for investigators seeking to adapt azosomes for photo-release of other compounds, as the physicochemical properties of the cargo, including its partitioning between the aqueous core and lipid bilayer, may influence encapsulation and release. A key component in our system was a commercially available optofluidic cannula, which allowed both delivering and measuring light using fiber photometry, and the ability to directly administer azosomes or other compounds into the brain region of interest. We initiated our studies with calcein-filled azosomes, and determined the stability, extent of leakage if any, spatial confinement around the site of infusion, and sensitivity to power and duration of the irradiation pulse. Some of these validation steps could be carried out *in vitro*, but we observed some differences in power sensitivity between *in vitro* and *in vivo* measurements (compare Fig. 2G and **Supplementary Fig. S3**). Based on our results, injection of 276 nL of azosomes through the cannula was sufficient to elicit measurable changes in local neuronal activity following photostimulation. We used a 1 s pulse at high power to mimic volume transmission, but shorter pulses can be used to deliver more transient signals. Compared to caged neuropeptide approaches^13^, which often require prolonged or repeated UV illumination to progressively uncage extracellular peptide pools, azosomes enable rapid release of the encapsulated neuropeptide cargo within seconds following brief irradiation pulses. This feature may better approximate the fast dynamics of endogenous neuropeptide signaling and provides greater temporal flexibility for studying neuromodulatory processes *in vivo*.

After validating that dye-filled azosomes work *in vivo* and determining the optimal power and duration for photo-releasing, we next characterized the properties of neuropeptide-filled azosomes. To validate the release of the neuropeptide *in vivo*, it is essential to measure the release, such as with a genetically encoded neuropeptide sensor. We used the OT sensor MTRIA_OT_ but first characterized the response of the sensor *in vivo* with exogenous neuropeptide. We examined the response to specific concentrations of peptide. Surprisingly, the sensitivity of MTRIA_OT_ appeared lower *in vivo*; that is the EC_50_ for MTRIA_OT_ is 20.5 nM^19^, but we found a minimum of 10 µM was needed to induce a MTRIA_OT_ response. This difference may be due to the large and rapid dilution of peptide inject *in vivo*. Importantly, the OT receptor antagonist OVTA completely occluded OT activation of MTRIA_OT_. We then used MTRIA_OT_ to detect the photostimulated release of OT from OT-azosomes. Photo-release of OT produced a sustained activation of MTRIA_OT_ that was inhibited by OVTA.

Having validated photo-release of OT *in vivo*, we next examined the effect of photoreleasing OT in the CA2 region of hippocampus just prior to a social behavior interaction test. In this experiment, we could administer the OT azosomes and directly control the timing of neuropeptide availability in the brain, overcoming limitations of existing approaches that rely on manipulation of neuropeptide-releasing neurons. In addition to observing the social behavior, we simultaneously measured OT levels using MTRIA_OT_ or neuronal activity using GCaMP. At the behavioral level, photo-releasing OT immediately before the gate lifted resulted in a reduced latency to initiate social interaction and shorter interaction duration following photorelease. Importantly, these changes occurred without altering the overall number of social-contact events, suggesting that temporally controlled OT delivery modulates specific components of social behavior rather than globally increasing social drive.

Most previous studies examining oxytocin signaling in hippocampal-associated social behaviors have focused on social memory and social recognition paradigms, typically using genetic or pharmacological manipulations during the acquisition phase of repeated social encounters^21,22^. In contrast, our experiments were designed to examine the acute behavioral consequences of temporally precise oxytocin release immediately prior to social interaction, enabled by the rapid kinetics of azosome-mediated photo-release. This temporal precision was incorporated with a custom-built behavioral platform that enabled subsecond detection of individual social-contact events, allowing precise alignment of oxytocin release with behavioral responses. Although we did not directly assess social memory, it remains possible that acute OT release in CA2 modulated familiarity-related processing, social salience, or stress-related responses associated with the encounter. Consistent with this possibility, altered hippocampal activity has been reported in oxytocin-deficient mice during social exposure^21^, and hippocampal circuits are also broadly implicated in stress and emotional processing^23^. Importantly, photo-release of OT reduced the latency to initiate interaction and shortened interaction duration without altering the total number of social-contact events, suggesting that temporally controlled OT signaling may modulate specific aspects of social processing or behavioral state rather than globally increasing social drive. Future studies will be required to determine whether azosome-mediated OT release also influences social memory formation or retrieval.

In addition to changes in behavior, photo-released OT altered the activity of CA2 glutamate neurons as well as modified the dynamics of endogenous OT during social interaction. In the absence of OT release, vGLUT1 neurons decreased in firing around social contact with a rebound of activity ∼ 10 s later. Similar changes in OT levels were measured with MTRIA_OT_, suggesting a close coupling between endogenous OT signaling and CA2 excitatory activity. Photo-release of OT produced a marked suppression of glutamatergic activity, as evidenced by reduced frequency and amplitude of calcium transients during social interaction. Photoreleased OT also reshaped endogenous OT signaling dynamics, attenuating the characteristic pre-contact decrease and post-contact increase observed under control conditions. Disruption of this coordinated pattern following photorelease indicates that elevating OT levels can alter circuit function in a manner that diverges from endogenous signaling regimes. Together with the behavioral results, these findings suggest a causal link between neuropeptide release and modulation of circuit activity and behavioral output.

Several mechanisms may account for the observed circuit effects. Previous studies have shown that OXTR activation can directly depolarize CA2 pyramidal neurons and promote burst firing^24^, consistent with a model of first-order neuromodulation in which OT enhances excitatory activity^25^. This framework aligns with the coordinated OT and vGLUT1 neuron dynamics observed under control conditions. In contrast, the suppression of excitatory activity following photorelease of OT suggests engagement of additional network mechanisms. One possibility is that elevated OT levels recruit local inhibitory interneurons, leading to a net reduction in excitatory output, a form of second-order neuromodulation described in other hippocampal subregions^26,27^. Alternatively, receptor saturation or desensitization may contribute to the attenuation of both OT signaling dynamics and neuronal activity. While the present data do not distinguish between these possibilities, they highlight how precise control over neuropeptide levels can reveal distinct regimes of circuit modulation that are not accessible through endogenous release alone.

Our results provide insight into the temporal dynamics of neuropeptide signaling *in vivo*. This observation is consistent with recent studies using genetically encoded sensors, which have revealed rapid components of neuropeptide dynamics in multiple brain regions. The ability to both measure and manipulate neuropeptide signaling with this level of precision highlights the importance of controlled delivery for understanding neuromodulatory function. Importantly, this approach provides a framework to causally link neuromodulator dynamics to circuit activity and behavior *in vivo*. The mechanisms underlying the rapid decrease and increase in neuropeptide levels, however, remain to be determined.

Azosomes offer several advantages over existing strategies for manipulating neuroactive molecules *in vivo*. Unlike optogenetic or chemogenetic approaches, which rely on endogenous release mechanisms and are subject to co-transmission of multiple signaling molecules, azosomes enable direct delivery of defined compounds. Compared to caged compounds, which require molecule-specific chemical modification and may present challenges in ensuring complete inactivity prior to uncaging as well as rapid peptide degradation *in vivo*, azosomes provide a formulation-based approach that is readily adaptable to a wide range of cargos. Importantly, the photoswitchable properties of Azo-PC lipids enable reversible trans-cis isomerization of the membrane, providing optical control over vesicle permeability and the temporal window of cargo release. Although cargo release itself is irreversible, subsequent illumination with 455 nm light can return Azo-PC to the trans configuration, reducing membrane permeability and thereby terminating further light-induced release, as previously demonstrated^16^. This reversible photoswitching provides an opportunity to regulate the duration of release and may enable dose-controlled delivery of bioactive molecules in future physiological and behavioral studies. We note some potential limitations. First, the azosome preparation contains a small fraction of free (non-encapsulated) OT, which could contribute to baseline effects following infusion. To mitigate this, we incorporated a delay between infusion and photostimulation to allow diffusion of free neuropeptide before triggering release, thereby improving separation between low- and high-OT conditions. Second, delivery was performed unilaterally and within a confined volume, which may limit the spatial extent of modulation and influence network-level interpretations; bilateral delivery could further enhance these effects and provide a more physiologically relevant approach.

In summary, this study establishes azosomes as a platform for temporally precise manipulation of neuropeptide signaling *in vivo* and demonstrates their utility for linking molecular, circuit, and behavioral processes. By enabling direct, externally controlled delivery of bioactive molecules with high temporal resolution, this approach provides a powerful framework for dissecting neuromodulatory mechanisms and can be readily extended to other neuropeptides, pharmacological agents, and brain regions.

## Materials and methods

### Calcein and Oxytocin loaded azosomes preparation

DSPC (>99%), 18:0-azo PC (830.1 g/mol, >99%), and cholesterol derived from ovine wool (>98%) were obtained from Avanti Polar Lipids Inc. Calcein sodium salt (sodium salt, 644.5 g/mol; Alfa Aesar, Cat. No. 108750-13-6) and Oxytocin (acetate salt, 1007.2 g/mol; Bachem, Product No. 4016373) were used in this study. All other reagents were of analytical grade and used as received. Azo-PC–based liposomes were prepared using the thin-film hydration technique as previously reported^28,29^. Briefly, DSPC, cholesterol, and azo-PC were dissolved in chloroform at a molar ratio of 58:30:12. The solvent was evaporated under a nitrogen stream, followed by overnight drying under vacuum. The resulting lipid film was hydrated with 1 mL of phosphate-buffered saline (PBS) containing either Calcein (75 mM) or Oxytocin (4 mM) at 65 °C for 1 h. The suspension was subjected to five freeze–thaw cycles (1 min in liquid nitrogen and 2 min in a 65 °C water bath) and then extruded 11 times through 400 nm and 200 nm polycarbonate membranes (Whatman, USA) using a Mini Extruder (Avanti Polar Lipids, USA). Unencapsulated calcein or oxytocin was removed by size-exclusion chromatography using a Sephacryl® S-500 HR column (Sigma-Aldrich, Cat. No. GE17-0613-10). Calcein-loaded azosomes were used immediately, whereas oxytocin-loaded azosomes were further purified by dialysis (Spectra Dialysis Tubing, 6–8 kDa MWCO). The hydrodynamic diameter and size distribution of the liposomes were determined by dynamic light scattering (Malvern Zetasizer Nano ZS) at room temperature.

Calcein Azosomes quantification: Calcein azosomes were irradiated with 365 nm light using a UV-LED light source (LC-L1V5, Hamamatsu, Japan) at various pulse durations (0.1–10 s) and power densities (175–1750 mW/cm²). For each condition, 100 µL of sample was added to a 96-well plate and irradiated with the LED light. Fluorescence intensity was then measured using a plate reader (excitation: 480 nm; emission: 520 nm). The release percentage was calculated by normalizing the fluorescence intensity of each condition to the maximum release obtained after 10 s of irradiation (**Supplementary Fig. S3C**).

### LC-MS Quantification of Oxytocin azosomes

200 µL of Oxytocin azosomes were irradiated with 365 nm light using a UV-LED light source (LC-L1V5, Hamamatsu, Japan) at various pulse durations (0.1–10 s) and power densities (175–1750 mW/cm²). The irradiated sample was filtered through a 0.1 µm filter to purify it prior to quantification. The release percentage was calculated by normalizing the fluorescence intensity of each condition to the maximum release obtained after 10 s of irradiation. The release profile of Oxytocin following irradiation was analyzed using LC-MS. Briefly, 5 μL of the aliquot was injected into the LC-MS system, separated with an eluent of water-acetonitrile containing 0.01% trifluoroacetic acid, starting from 100% water to a 50% water-acetonitrile mixture over 7 minutes. The Oxytocin peak was measured based on its height, and its identity was confirmed by mass spectrometry, corresponding to a molecular weight of 1007.3 kDa.

### Azosomes Cryo-electron microscopy imaging

3.5 µL of azosomes was applied to a lacey carbon 300-mesh copper grid (Ted Pella) that was pre-treated by 80 s glow discharge at 30 mA in a PELCO easiGlow glow discharge apparatus (Ted Pella). After a 3 s wait, the grid was blotted for 3.5 sec and plunge-frozen in liquid ethane using Vitrobot Mark IV (Thermo Fisher Scientific) at 95% humidity and 7° C. Grids were imaged on a 200 kV Glacios microscope (Thermo Fisher Scientific) at the Cryo-Electron Microscopy Facility (CEMF) at UT Southwestern Medical Center with a K3 direct detection camera (Gatan). Movies were acquired using SerialEM ver 4.2.0 beta at 45,000 X magnification (pixel size of 0.874 Å) for 2.1 secs at dose rate of 18 e^−^/pixel/s in non-CDS mode, with an accumulated total dose of 50 e^−^/Å^2^ over 50 frames at a defocus of −4 μm. Movies were aligned in SerialEM and images were saved as jpg.

### Mice

All experiments were conducted in accordance with institutional guidelines and an approved protocol by the Institutional Animal Care and Use Committee (IACUC) at the Icahn School of Medicine at Mount Sinai. For all experiments described below, male C57BL/6J mice (3–6 months old, approximately 20–35 g; Jackson Laboratory) were used. Mice were housed in groups of 3–5 per cage under a 12-hour light/dark cycle in a temperature- and humidity-controlled environment, with ad libitum access to food and water. Littermates of the same sex were randomly assigned to experimental groups.

### Stereotaxic infusion of liposomes and azosomes

Adult mice were anesthetized with isoflurane in oxygen and placed in a stereotaxic frame (5% for induction, 2% for maintenance; 1–3 L/min). Bilateral craniotomies were performed above the nucleus accumbens core (NAc core) at the following coordinates relative to bregma: AP +1.0 mm, ML ±1.3 mm. Glass micropipettes were prepared from fire-polished capillaries (WPI; 1.14 mm outer diameter, 3.5ʺ length; catalog #504949) using a micropipette puller (Narishige PC-10; heater setting 58.9) to produce a long, narrow shank. The tip was subsequently trimmed with fine scissors to achieve an opening diameter of 7–9 μm, following established guidelines. Micropipettes were backfilled with mineral oil and front-loaded with 2–4 μL of calcein-loaded liposome or azosome suspension. The pipette was lowered to the target depth (DV −4.5 mm), and injections were performed using a nanoliter injector (Nanoliter 2010, WPI). A total volume of 220.8 nL was delivered as 16 discrete injections of 13.8 nL over 4 minutes (injection rate: 55.2 nL/min). Following infusion, the pipette was left in place for 5 minutes to allow diffusion and minimize reflux, and then slowly withdrawn to prevent backflow along the injection tract.

### Dual fiber-optic cannula implantation and photometry setup

Following infusion, dual fiber-optic cannula was implanted to enable simultaneous fluorescence recording and light delivery. The cannula (Doric Lenses; TFC_400/430-0.48_5mm_TSM2.6_FLT), with a 400 μm core diameter optical fiber, was positioned bilaterally at the same AP and ML coordinates as the injection sites (NAc core, relative to bregma: AP +1.0 mm, ML ±1.3 mm). It was then lowered slowly (0.5–1.0 mm/min) while monitoring fluorescence signals in real time using the photometry system. Mono fiber-optic patch cords (Doric; MFP 400/430/1100-0.57, 3 m, low autofluorescence) were used for signal collection and light delivery. As the fiber tips approached the injection sites, fluorescence increased, confirming proximity to the vesicle-containing region (**Fig. 1H i**) at a depth of DV −4.5 mm. Once optimal positioning was achieved, cannulas were secured to the skull using dental cement (C&B Metabond®, Parkell; radiopaque tooth shade powder, quick base, and universal catalyst components). Fluorescence signals were recorded during fiber approach and following light stimulation. Light pulses were delivered at 175 mW/cm² with durations of 200 ms or 2 s. Fluorescence increases reflect release of calcein from vesicles due to reduced self-quenching upon dilution in the extracellular space.

### Fiber photometry acquisition

Fiber photometry recordings were performed using two optical configurations depending on the experimental application. In both cases signals were recorded at 1017.25 Hz using Synapse software.

For experiments involving UV-triggered azosome photorelease (calcein, MTRIA-OT, and jGCaMP7s), fluorescence was acquired using a TDT RZ5P processor (Tucker-Davis Technologies). Green fluorescence was excited using a 490 nm LED (M490F2, Thorlabs) and collected through an excitation filter (MF469-35), long-pass dichroic mirror (DMLP425R), dichroic beamsplitter (MD498), and emission filter (MF525-39) (Thorlabs). Azosome photorelease was achieved by replacing the standard 405 nm LED (M405F1) with a 365 nm LED (M365FP1, Thorlabs) while maintaining the same optical path, allowing simultaneous fluorescence recording and UV stimulation through a single optical fiber.

For experiments monitoring Calcein Red™ (sodium salt, 780.7 g/mol; Item No. 20633) during infusion of free OT or OT-loaded azosomes, fluorescence was acquired using a three-color fiber photometry system consisting of a TDT RZ10X processor equipped with integrated 405 nm (Lx405), 465 nm (Lx465), and 560 nm (Lx560) LEDs (Tucker-Davis Technologies). Green and red fluorescence were separated using a Doric six-port Fluorescence MiniCube (FMC6_IE(400–410)_E1(460–490)_F1(500–540)_E2(555–570)_F2(580–680)_S, version 2017), enabling simultaneous detection of green and red fluorophores. Because this configuration did not include a 365 nm excitation source, it was used exclusively for fluorescence monitoring and not for photo-triggered azosome release.

Optical power at the fiber tip was measured before each experiment using a PM100D Compact Power and Energy Meter coupled to an S150C silicon photodiode power sensor (350–1100 nm; Thorlabs). Power density (mW/mm²) was calculated from the measured output power and the cross-sectional area of the optical fiber.

### Fiber photometry analysis

Fluorescence traces were analyzed in Python as changes in fluorescence over time. For calcein-loaded azosomes, increases in fluorescence reflected cargo release, as dilution of encapsulated calcein following release reduced self-quenching and increased fluorescence intensity^18^. Fluorescence traces were aligned to the onset of UV stimulation or infusion events, and signals were expressed as ΔF/F (or z-score, depending on your analysis pipeline) for visualization and comparison across experiments.

### Histology

At the end of experiments, mice were deeply anesthetized with ketamine/xylazine (e.g., 100/10 mg/kg, i.p.) and transcardially perfused with phosphate-buffered saline (PBS), followed by 4% paraformaldehyde (PFA) in PBS. Brains were rapidly extracted, post-fixed overnight in 4% PFA at 4°C under gentle agitation, and then rinsed in PBS. Brains were embedded in 5% agarose (in PBS) and sectioned into 40-μm coronal slices using a vibratome. Sections were stored in PBS containing 0.02% (w/v) sodium azide until further processing. For nuclear staining, sections were incubated for 1 hour at room temperature in the dark with DAPI (0.1 mg/mL; Sigma-Aldrich, Cat# D9542), followed by three washes in PBS (5 minutes each). Sections were mounted onto glass slides, air-dried for 15–30 minutes (protected from light), and coverslips mounted with using Fluoromount-G (Thermo Fisher Scientific, Cat# 5018788). Slides were sealed with nail polish. Whole-slide images were acquired using a NanoZoomer S60 Digital Slide Scanner (Hamamatsu) to verify injection sites and fiber placement within the hippocampal CA2 and DMS (**Supplementary Fig. S4**).

### Agarose phantom preparation and opto-fluidic imaging

To model infusion and light-triggered release under tissue-like mechanical conditions, opto-fluidic experiments were performed in a 0.5% (w/v) agarose gel prepared in phosphate-buffered saline (PBS). This low-density agarose matrix provides a porous environment that approximates the mechanical properties of brain tissue and is commonly used to assess diffusion and dispersion of injected solutions. Opto-fluidic cannulas were inserted into the agarose block, and calcein-loaded azosomes were infused through the fluid injector. Fluorescence imaging was performed using a stereo fluorescence microscope (Olympus MVX10) equipped with a mercury lamp light source and standard GFP filter sets (excitation ∼470–490 nm, emission ∼510–550 nm), suitable for detection of calcein fluorescence. Time-lapse images were acquired during both infusion and subsequent photostimulation. For photorelease experiments, 365 nm light was delivered through the optical fiber of the cannula at 175 mW/cm² for 30 s. Fluorescence changes were monitored in real time, capturing both the initial distribution of vesicles within the agarose matrix and the subsequent increase in fluorescence associated with light-triggered release. Fluorescence intensity was quantified from regions of interest (ROIs) centered around the infusion site. Changes in fluorescence were expressed as ΔF/F relative to baseline fluorescence prior to photostimulation.

### Opto-fluidic cannula implantation

Opto-fluidic cannulas (Doric Lenses; OmFC ZF1.25 400/430-0.48 3 mm FLT) consist of a 400 μm core diameter optical fiber integrated with a fluid-guiding channel, together forming an elliptical cross-section of 450 × 650 μm. Cannulas were stereotaxically implanted and secured to the skull using dental cement as described above. For experiments targeting the hippocampus, cannulas were implanted into the right CA2 region (AP −2.3 mm, ML −2.4 mm, DV −2.1 mm relative to bregma). For experiments targeting the dorsomedial striatum (DMS), cannulas were implanted at AP +0.5 mm, ML −1.3 mm, DV −2.6 mm. Special attention was given to the orientation of the opto-fluidic cannula to ensure appropriate positioning of the optical fiber and fluid delivery channels. In DMS experiments, the cannula was implanted such that the fluid outlet was oriented laterally, away from the lateral ventricle (LV), to prevent unintended fluid entry into the ventricular system. Improper positioning of the fluid channel in close proximity to the LV can result in backflow or cerebrospinal fluid (CSF) influx upon removal of the cannula plug, compromising infusion stability. To avoid this, the orientation of the cannula was adjusted depending on the target hemisphere, ensuring that the fluid delivery port was consistently positioned away from ventricular structures. Mice were allowed to recover for 2–3 weeks prior to experimentation.

### Fluid injector assembly and *in vivo* opto-fluidic infusion and photostimulation

To enable controlled infusion via the opto-fluidic cannula, a custom injection assembly was constructed by coupling a pulled glass micropipette to polyethylene tubing (Doric; F220-0505-2) and the cannula fluid injector (Doric; FI_OmFC-ZF_100/170_3.1 mm). Glass micropipettes were pulled as described above and the tip trimmed to an aperture of 40–60 μm to reduce flow resistance and facilitate stable delivery. The micropipette tip was connected to the tubing and sealed using a minimal amount of heated adhesive, ensuring both mechanical stability and a tight junction. The distal end of the tubing (10-20 cm length) was connected to the fluid injector via a stainless steel inlet (25G), forming a continuous fluidic pathway. Prior to each experiment, the integrity of the assembly was verified by applying pressure using a syringe to confirm unobstructed flow and absence of leaks, with particular attention to the glass–tubing interface. This quality control step was critical to ensure reliable and reproducible infusion.

For experimental sessions, mice were anesthetized with isoflurane as described above and connected to the fiber photometry setup. Optical connections were established using mono fiber-optic patch cords (Doric; MFP 400/430/1100-0.57) coupled to the implanted cannula via Ø1.25 mm quick-release interconnects (Thorlabs; ADAL3), ensuring stable coupling and minimizing motion-related noise. The custom injection assembly was prepared and filled with mineral oil and front-loading ≥4 μL of calcein-loaded azosome suspension. The cannula plug (Doric; PLG_OmFC_195_3.1) was then carefully removed and replaced with the fluid injector connected to the assembly. For each recording session, azosomes were delivered as two consecutive injections of 32 nL each, separated by 15 s (total volume: 64 nL). Fluorescence signals were continuously recorded throughout the experiment. Light stimulation was delivered 75 s after the start of infusion to allow stabilization of the fluorescence signal. Photostimulation was performed using 365 nm light delivered through the optical fiber of the cannula. Two power conditions were tested: low power (0.22 mW at fiber tip; 175 mW/cm²) and high power (2.20 mW; 1750 mW/cm²). Light pulse durations ranged from 1 ms to 5 s (1, 2, 5, 10, 25, 50, 100, 200, 500 ms, and 1, 2, 5 s), with each condition tested in independent trials.

For stability experiments, photostimulation parameters were fixed to 1 s pulses at 365 nm delivered at high power, and the interval between azosome infusion and photostimulation was systematically varied. Photorelease was triggered at time points ranging from 1 min to 14 h post-infusion (1, 5, 15, 30, 45, 60, 120 min, and 3, 4, 8, and 14 h). For short time intervals (≤1 h), mice were maintained under isoflurane anesthesia throughout the experiment. For longer intervals (>1 h), mice were returned to their home cage following infusion and subsequently re-anesthetized prior to photostimulation and recording to minimize prolonged exposure to isoflurane.

Fluorescence traces were analyzed as ΔF/F, using a 30 s baseline period prior to light stimulation. To enable comparison across sessions and animals, ΔF/F traces were further normalized within each recording session to the maximal response obtained with the longest pulse duration (5 s) and expressed as a percentage. Peak fluorescence, time to peak, and decay were extracted from individual traces. Peak fluorescence was defined as the maximum ΔF/F value following stimulation. Time to peak was calculated as the time from stimulation onset to peak fluorescence. Decay was defined as the percentage decrease in signal between 25 s and 115 s post-stimulation. Values exceeding 100% were occasionally observed, likely reflecting partial photobleaching at longer stimulation durations rather than increased cargo release.

### Viral delivery and *in vivo* oxytocin detection

For detecting OT, we used pAAV.Syn.MTRIA-OT, which was a gift from Hiroshi Hibino & Daisuke Ino (Addgene # 184594; RRID: Addgene_184594). AAV2/8-Syn.MTRIA_OT_ -WPRE (4.74 × 10¹³ GC/mL was prepared at Harvard Boston Children’s Hospital viral core. For oxytocin sensing experiments, opto-fluidic cannulas were implanted in the dorsomedial striatum (DMS) as described above. After a 2–3 week recovery period, pAAV.Syn.MTRIA-OT was infused through the cannula. Viral solutions were diluted (3:4) and supplemented with calcein (final concentration 25 μM) to allow real-time monitoring of infusion cannula (**see Fig. 4**). Animals were allowed 3–4 weeks for sensor expression before recordings. During experimental sessions, mice were connected to a three-color fiber photometry system (TDT RZ10x) equipped with excitation sources at 405 nm (isosbestic), 465 nm (MTRIA_OT_ ), and 560 nm (Calcein Red), and a fluorescence minicube (Doric; FMC6_IE(400–410)_E1(460–490)_F1(500–540)_E2(555–570)_F2(580–680)_S) enabling simultaneous detection of green and red emission channels. Oxytocin solutions were prepared at increasing concentrations (10 nM to 100 μM) in vehicle containing Calcein Red (10 μM final concentration) to monitor infusion. For each condition, solutions were delivered through the opto-fluidic cannula using the injection parameters described above. For pharmacological validation, OT (10 μM) and the oxytocin receptor antagonist OVTA (100 μM) were sequentially infused within the same session, with a 15 min interval between infusions. In combined conditions, OT was delivered in the presence of OVTA to assess receptor-dependent blockade of the sensor response. Fluorescence signals were processed as described above.

### *In vivo* photorelease of OT-loaded azosomes

Experiments were performed in mice expressing MTRIA_OT_ in the DMS and implanted with opto-fluidic cannulas as described above. All recordings were conducted under isoflurane anesthesia in a stereotaxic frame. For each experimental session, different solutions (vehicle, OT-azosomes, OT, or OVTA) were sequentially delivered through the opto-fluidic cannula using the custom injection assembly. Each cycle consisted of: (i) loading the injector with the selected solution containing 10 μM Calcein Red, (ii) insertion into the cannula, (iii) a 1–2 min stabilization period, (iv) infusion (4-8 × 13.2 nL, 15 s interval), and (v) light stimulation delivered 2–5 min after infusion. Sequential conditions were separated by 15 min. Photostimulation was delivered using 365 nm light through the optical fiber of the cannula. For OT-azosome experiments, a single high-power pulse (5 s; 1750 mW/cm²) was used. In OVTA experiments, a 1 s pulse followed by a 5 s pulse (2 min later) was applied to assess release completeness. Short (1 s) pulses were sufficient to elicit maximal responses, indicating rapid depletion of releasable cargo. During infusion, signals were acquired using a 3-color photometry system enabling simultaneous recording of MTRIA_OT_ (green channel), Calcein Red (red channel), and isosbestic control. During photorelease, recordings were performed using a configuration compatible with UV stimulation (365 nm) while maintaining MTRIA_OT_ detection.

### Data analysis

Fluorescence signals were processed as ΔF/F₀ using a baseline defined prior to infusion or irradiation events. Infusion responses were quantified as peak ΔF/F₀ following delivery, whereas irradiation responses were quantified as peak changes following light stimulation, excluding transient optical artifacts.

### Social interaction assay with integrated opto-fluidic delivery and photometry recording

Mice were implanted with opto-fluidic cannulas targeting the hippocampal CA2 region and infused with AAV encoding MTRIA_OT_ (as described above) or, in a separate cohort, a Cre-dependent calcium indicator AAV9-Syn-FLEX-jGCaMP7s-WPRE (3.1 × 10¹³ GC/mL; Addgene #104491-AAV9) in Vglut1-Cre male mice. After 3–4 weeks of expression, experiments were conducted in male mice. For each session, OT-azosomes were infused under isoflurane anesthesia using the opto-fluidic system. The total infused volume (4 injections of 69 nL) was adjusted based on Calcein Red fluorescence to ensure comparable delivery across conditions within each animal. Following infusion, mice were returned to their home cage for 10 min to recover from anesthesia before behavioral testing. Animals were then placed in the isolated compartment of the behavioral arena and allowed to acclimate for an additional 10 min. During irradiation sessions, brief light pulse (365 nm, 1 s, high power) was delivered 4–5 min prior to gate opening, whereas control sessions were performed without irradiation. Social interaction was initiated by opening the gate, allowing access to a female stimulus enclosed in a barred cup. Behavioral sessions were performed in a within-subject design, with each animal tested under both conditions. Conditions were counterbalanced across sessions such that each animal alternated between irradiation and non-irradiation sessions.

Animal position was tracked using top-down video recording and analyzed using the Bonsai framework. Social-contact events were defined by the conjunction of three criteria: (i) gate open state, (ii) proximity to the stimulus (based on position tracking), and (iii) activation of a capacitive sensor integrated into the stimulus enclosure. The gate and capacitive sensing system were controlled by a custom Arduino-based board (**Supplementary Fig. S5**). The social stimulus enclosure (cup) was modified by lining the external bars with conductive copper tape, which was electrically connected via jumper wires to the microcontroller. This configuration allowed the cup to function as a capacitive sensor, such that physical contact from the test animal resulted in a measurable change in capacitance. Signal detection was implemented using a capacitive sensing approach based on the CapacitiveSensor library (Paul Badger), in which changes in charge–discharge timing are used to infer contact events. This setup enabled reliable detection of direct physical interaction with the enclosure while minimizing false positives from proximity alone.

Fluorescence signals (MTRIA_OT_ or vGLUT1-GCaMP) were recorded continuously and analyzed using custom Python-based pipelines. Signals were aligned to behavioral events, and social-contact epochs were identified based on the criteria described above to compute event-aligned averages and raster representations. Traces were z-scored using the mean and standard deviation calculated from the pre-irradiation baseline period (from session start to immediately before light delivery), and event-aligned traces were further baseline-normalized using a pre-contact window prior to each event. For MTRIA_OT_ signals, pre-contact trough amplitude and post-contact peak responses (5–15 s window) were quantified for each condition. For vGLUT1-GCaMP recordings, calcium transients were detected using peak-finding algorithms implemented in SciPy (scipy.signal.find_peaks), with thresholds for prominence, width, and inter-peak distance to identify physiologically relevant events. Transient frequency and amplitude (prominence) were quantified for each social-contact epoch and compared across conditions.

## Acknowledgements

We thank members of the Slesinger and Qin labs for helpful discussions. We are grateful to Joyce Fung for assistance with CryoEM imaging at the University of Texas Southwestern Medical Center and to Katarzyna Ciałowicz Farrugia and the Microscopy CoRE at the Icahn School of Medicine at Mount Sinai for microscopy support and technical assistance. This work was supported in part by grants from NSF (2123830) and NIH (R01NS110499). Biorender was used to generate schematics in Figures.

## Notes

### Competing Interest Statement

The authors have declared no competing interest.

