## Supplementary Figures for "PHOTORELEASE OF OXYTOCIN *IN VIVO* USING PHOTOSWITCHABLE NANOVESICLES MODULATES HIPPOCAMPAL CIRCUIT DYNAMICS AND SOCIAL BEHAVIOR"

**Supplementary Video 1**. Opto-fluidic infusion and photorelease of calcein-loaded azosomes in agarose.

**Supplementary Figure S1**. Area under the fluorescence response curve as a function of light pulse duration.

**Supplementary Figure S2**. *In vitro* characterization of oxytocin (OT) release from azosomes.

**Supplementary Figure S3**. Azosomes preparation workflow and *In vitro* characterization of calcein release from azosomes.

**Supplementary Figure S4**. Histological verification of opto-fluidic cannula placement.

**Supplementary Figure S5**. Design of the Arduino-based touch-sensor for automated social interaction.

#### Supplemental Figures

**Supplementary Video 1. Opto-fluidic infusion and photorelease of calcein-loaded azosomes in agarose.** Time-lapse fluorescence microscopy showing infusion of calcein-loaded azosomes through an opto-fluidic cannula into a 0.5% agarose gel, followed by 365 nm photostimulation. Two sequential infusion/photostimulation cycles are shown. Photostimulation induces calcein release from azosomes, resulting in fluorescence dequenching and subsequent diffusion from the infusion site. Related to Fig. 2A.

**Supplementary Figure S1. Area under the fluorescence response curve as a function of light pulse duration.** Area under the fluorescence response curve (AUC) as a function of light pulse duration at low and high power, corresponding to the experiments shown in **Figure 2G**. Curves were fitted using a standard four-parameter Hill equation. Data are presented as mean ± SEM (n = 4–10).

**Supplementary Figure S2. *In vitro* characterization of oxytocin (OT) release from azosomes**. (**A**) Schematic illustrating light-triggered release of encapsulated oxytocin from azosomes upon UV irradiation. (**B**) Stability of OT-loaded azosomes incubated at 37 °C for 2 h, showing minimal passive release (<5%) over time. (**C**) Quantification of OT released *in vitro* following UV irradiation with increasing light pulse durations (0, 0.5, 1, and 10 s). Bars represent the concentration of released OT (µM).

**Supplementary Figure S3. Azosomes preparation workflow and *In vitro* characterization of calcein release from azosomes.** (**A**) Schematic illustrating light-triggered release of encapsulated calcein from azosomes. (**B**) Percentage of calcein released as a function of light pulse duration (0.1, 0.5, 1, 2 and 5 s). under low- and high-power irradiation. Curves were fitted using a standard four-parameter Hill equation. (**C**) Schematic illustration of the azosome preparation process and subsequent removal of free cargo.

**Supplementary Figure S4. Histological verification of opto-fluidic cannula placement.** (**A**) Representative coronal brain sections showing opto-fluidic cannula implantation in the dorsomedial striatum (DMS). (**B**) Representative coronal brain sections showing opto-fluidic cannula implantation in the hippocampal CA2 region. The fiber tract is visible in each section, confirming accurate targeting of the intended brain regions. Two representative sections are shown for each target.

**Supplementary Figure S5. Design of the Arduino-based touch-sensor for automated social interaction.** (**A**) Schematic of the custom-built behavioral apparatus. The system consists of an Arduino-controlled servo gate that alternates between isolation and interaction periods, a capacitance touch sensor connected to copper tape surrounding the exterior of the stimulus cup to detect physical contact by the test mouse. **(B)** Representative synchronized video frames acquired from top and side views during isolation (gate closed) and social interaction (gate open). (**C**) Synchronized behavioral traces showing touch detection (blue), mouse x-position (yellow), and gate state (purple). See Methods for a detailed description of the apparatus and behavioral protocol.

### Figure S1

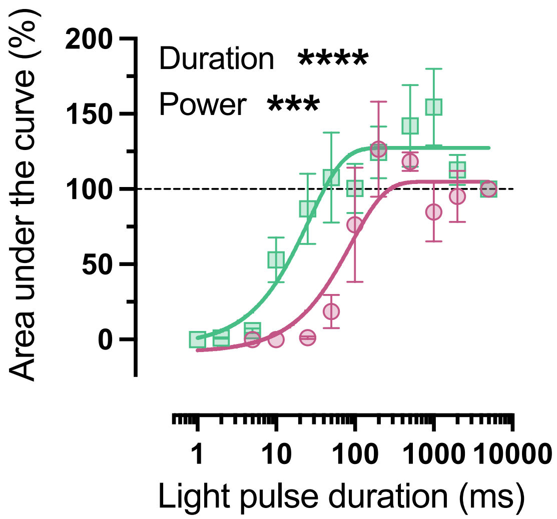

Area under the curve (%)
200
150
100
50
0
Duration ****
Power ***
1
10
100
1000
10000
Light pulse duration (ms)

### Figure S2

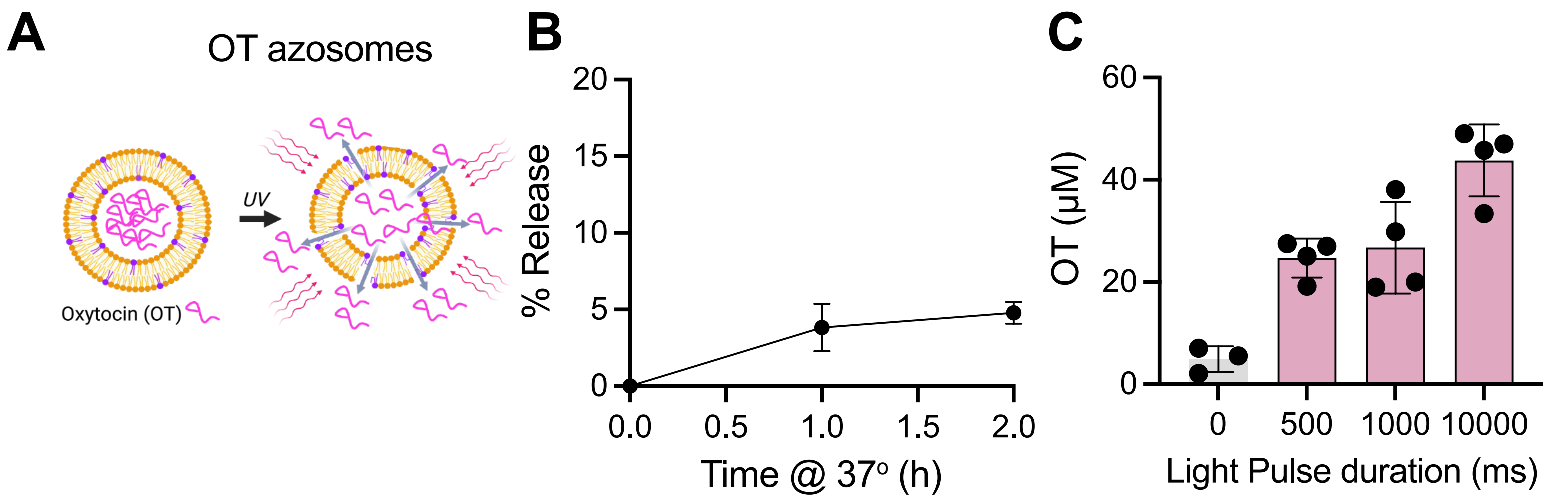
A
OT azosomes
UV
Oxytocin (OT)
B
% Release
20
15
10
5
0
0.0
0.5
1.0
1.5
2.0
Time @ 37° (h)
C
OT (μM)
60
40
20
0
0
500
1000
10000
Light Pulse duration (ms)

### Figure S3

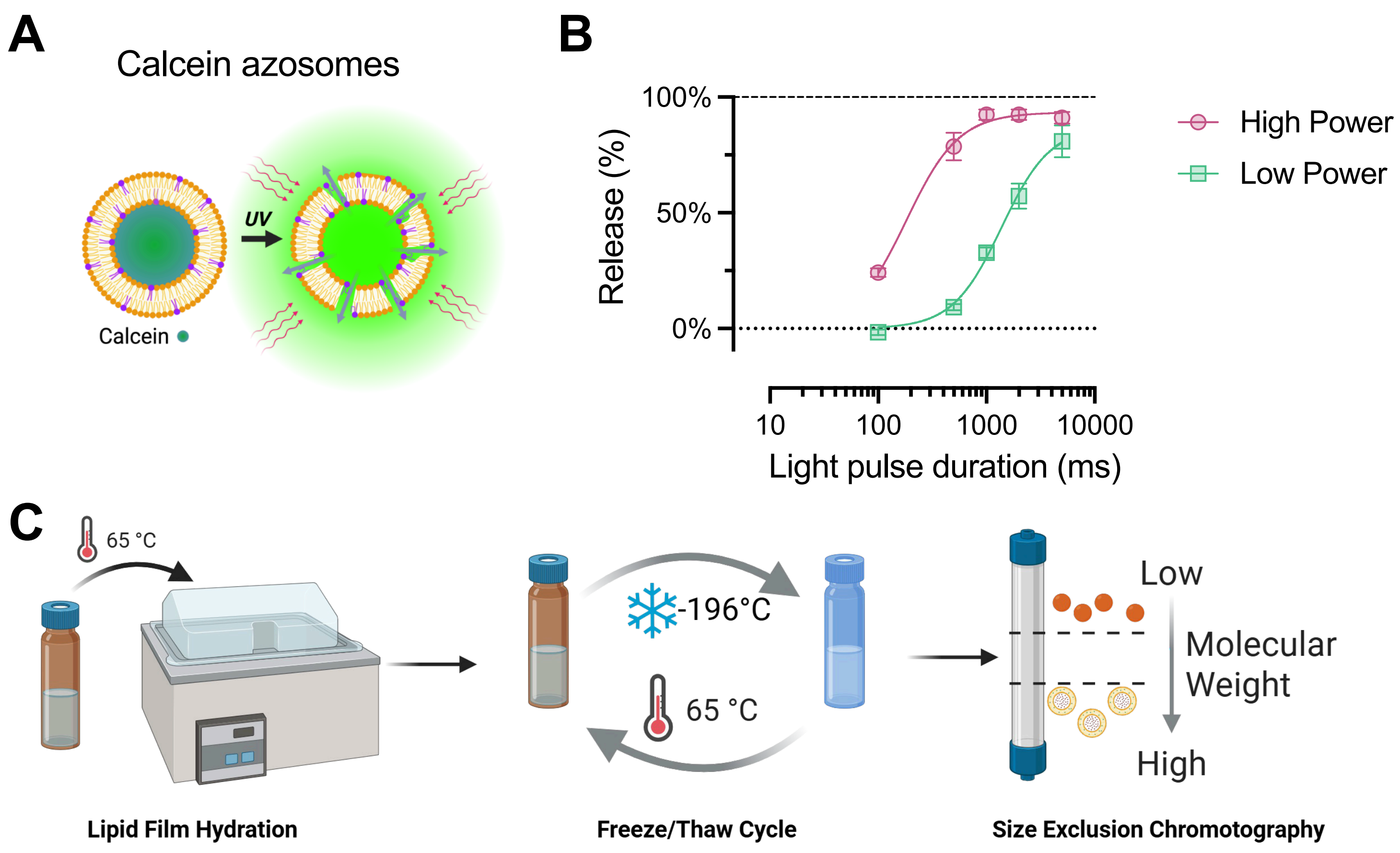


$$\textbf{\textit{Release}}\ (\%) = \frac{I_{Release} - I_{initial}}{I_{Total} - I_{initial}} \times 100\%$$

### Figure S4

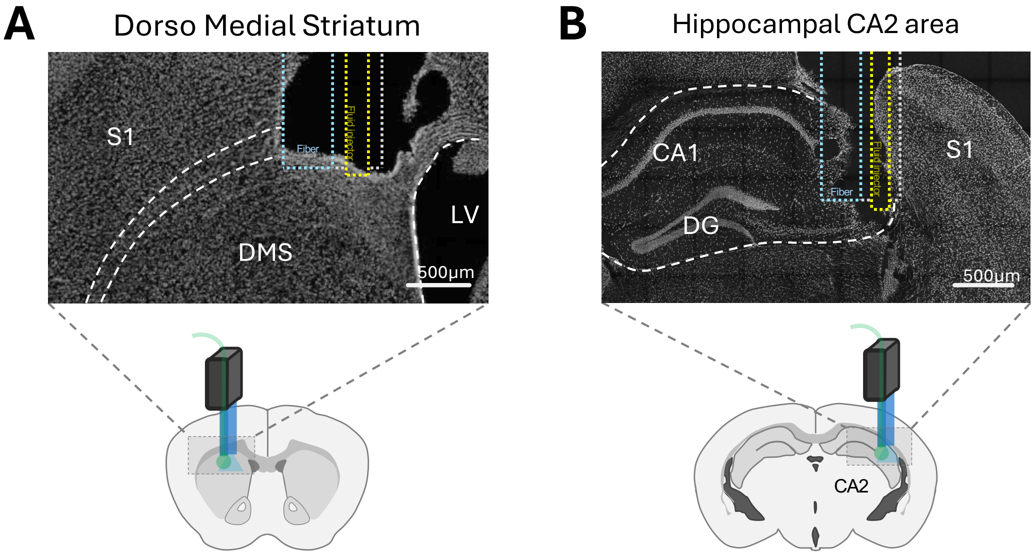
A
Dorso Medial Striatum
S1
Fiber
LV
DMS
500µm
B
Hippocampal CA2 area
CA1
S1
Fiber
DG
500µm
CA2

### Figure S5

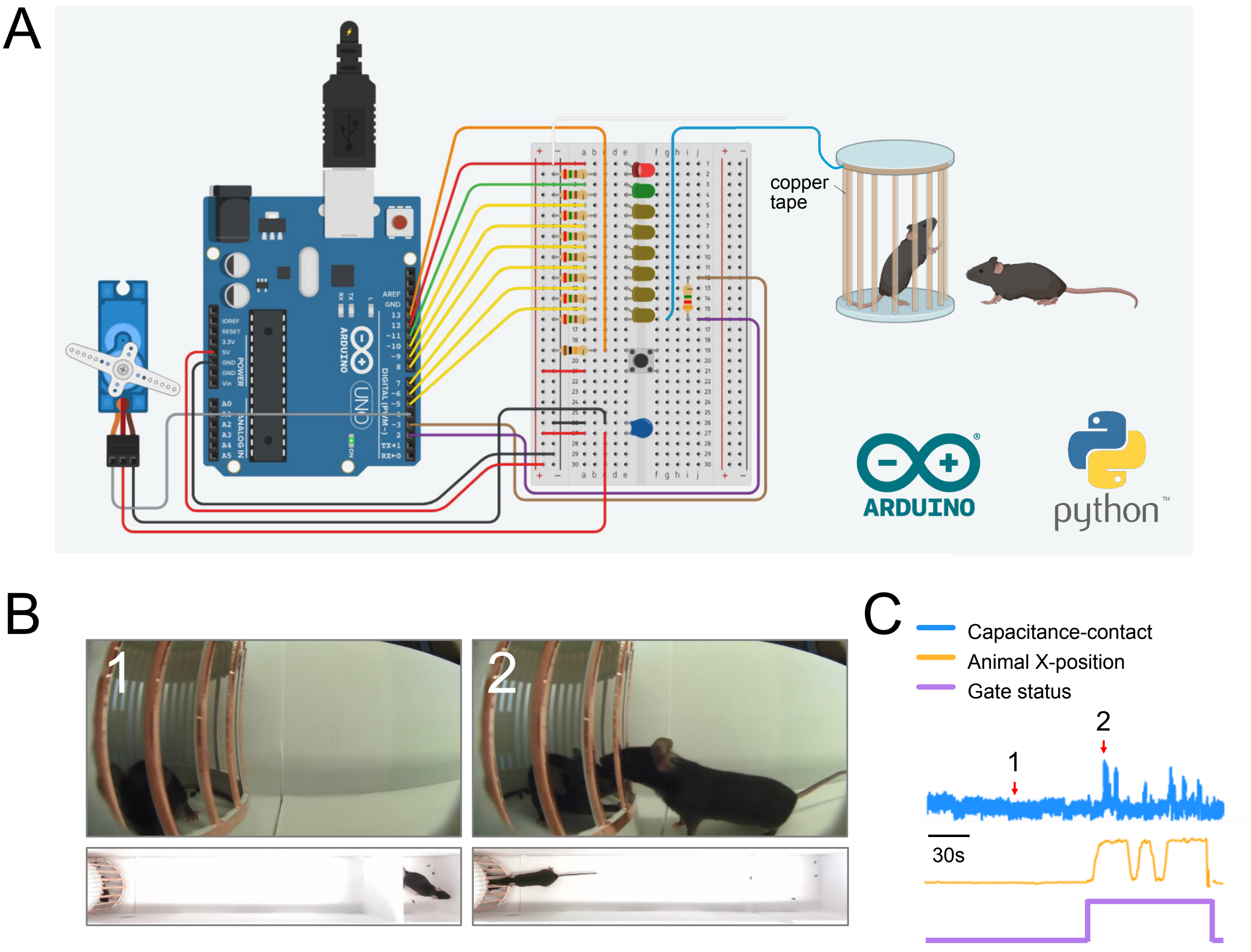
A
copper tape
ARDUINO
python
B
1
2
C
Capacitance-contact
Animal X-position
Gate status
1
2
30s
